# Efficient protein structure generation with sparse denoising models

**DOI:** 10.1101/2025.01.31.635780

**Authors:** Michael Jendrusch, Jan O. Korbel

**Affiliations:** European Molecular Biology Laboratory (EMBL), Genome Biology Unit, Heidelberg, Germany; Collaboration for joint PhD degree between EMBL and Heidelberg University, Faculty of Biosciences

**Keywords:** Computational protein design, denoising diffusion models

## Abstract

Generating designable protein backbones has become an integral part of machine learning-assisted approaches to protein design. Together with sequence design and structure predictor-based filtering, it forms the backbone of the computational protein design pipeline. However, current protein structure generators face important limitations for large proteins and require retraining for protein design tasks unseen during model training. To address the first issue, we introduce `salad`, a family of `s`parse `al`l-`a`tom `d`enoising models for protein backbone generation. Our models are notably faster than the state-of-the-art while matching or improving designability and diversity, and generate designable structures for protein lengths up to 1,000 amino acids. To address the second issue, we combine `salad` with structure-editing, a strategy for expanding the capability of protein denoising models to unseen tasks. We apply our approach to a variety of protein design tasks, from motif-scaffolding to multi-state protein design, demonstrating the flexibility of `salad` and structure-editing.

## I INTRODUCTION

Computational protein design aims to generate protein sequences and structures with specified folds, functions, and dynamics. Protein design tasks are varied, starting from designing proteins with specified secondary structure [1, 2] or symmetry [3–5] and producing scaffolds for known functional motifs [2, 6, 7] to designing potent binders for protein targets [1, 8–10] as well as multi-state protein sequences which can switch folds under different conditions [11–13]. These design tasks enable powerful applications in basic research and industry, for example designing or optimizing enzymes [14–16], antibodies [8], vaccine scaffolds [17, 18] and biosensors [12]. Many recent approaches to *de novo* protein design follow the same general pipeline of candidate backbone generation, followed by sequence design and filtering for likely successful designs using a protein structure predictor [1, 2, 19, 20] followed by experimental testing. This approach is enabled by the introduction of powerful protein structure predictors such as AlphaFold 2 [21] and ESMfold [22] combined with ProteinMPNN [23] as the *de facto* standard way of designing amino acid sequences given a backbone.

While sequence design and structure prediction are mostly standardized, a multitude of different approaches fill the role of protein backbone generation. Knowledge-based design coupled with backbone generation using Rosetta [24] plays a leading role especially for complex protein design tasks, such as enzyme design [15, 16], multi-state protein design [11, 12] and the design of proteins with strong geometric and sequence constraints [3, 4]. These methods are supported by machine learning based approaches, which are capable of solving simpler design tasks without relying on Rosetta or manual design: Protein structure hallucination methods [10, 19, 25–28] invert structure predictors using search or gradient descent to generate sequences with high-confidence predicted structures. These sequences are often adversarial and therefore discarded in favour of ProteinMPNN sequence designs [10, 19, 25]. Protein denoising diffusion models (DDPM) [29] iteratively generate proteins from random noise by learning to remove noise from corrupted protein sequences [13, 30] or structures [1, 2, 20, 31, 32]. Diffusion models have a runtime advantage over hallucination-based methods as they do not require optimization over a structure predictor [1, 19]. Recently, protein diffusion models have been successfully applied to solve various protein design tasks from unconstrained *de novo* protein design [1, 2, 20] to protein binders and complexes [1, 20].

While current protein diffusion models have shown impressive performance for small protein generation, their performance deteriorates with protein sequence length *N* [1, 2, 20, 31]. Excluding Chroma [20], the majority of protein diffusion models use architectures derived from either AlphaFold 2 [21] or RoseTTAfold [33], inheriting their runtime complexity. The use of amino acid pair features introduces a lower complexity bound of *O*(*N*^2^) and pair attention mechanisms increase this complexity to *O*(*N*^3^) [21, 33]. Along with decreased runtime performance, these models also experience a drop in designability with increasing *N*. To the best of our knowledge no protein structure diffusion model has reached the designability of hallucination-based approaches on large proteins [19]. While protein backbone hallucination enables large protein design, this comes at a significant runtime cost compared to diffusion models [19]. This greatly reduces throughput and may limit the applicability of backbone hallucination to large protein design tasks with lower per-design success-rates compared to unconditional monomer design.

Another issue with current protein structure diffusion models is the need for additional training to solve specific protein design tasks. RFdiffusion and Genie are separately trained with protein motif conditioning to scaffold functional protein motifs [1, 2]. While Chroma’s conditioners allow for training-free adaptation of the model for different tasks, implementing new conditioners is not straight forward and requires the development of custom energy functions [20]. Thus, there is a significant barrier to applying existing diffusion models to novel design tasks.

To address these issues, we introduce `salad` (`s` parse`al`l-`a`tom`d` enoising), a family of efficient protein generative models with sub-quadratic complexity. We train our models with a denoising diffusion objective [1, 29] to remove noise from corrupted protein backbones. Starting from a sparse transformer architecture [20, 34, 35], we investigate the impact of different model features and noise schedules on the designability and diversity of generated proteins. We find that our models are capable of generating diverse and designable backbones for proteins up to 1,000 amino acids long. `salad` matches or outperforms state-of-the-art diffusion models [1, 2] in terms of designability while drastically reducing runtime. By editing the outputs of our models at each denoising step, we enable rapid prototyping of protein design unseen during training. Combining `salad` with structure-editing, we tackle a variety of protein design tasks, generating designable backbones with specified shapes [20], scaffolds for functional protein motifs [1, 2, 6], repeat proteins [3, 4, 13] and multi-state proteins that adopt distinct folds when cleaved [13]. This way `salad` can be used as an efficient plug-and-play replacement for other backbone generators in existing protein design pipelines.

## II. RESULTS

### A. Sparse protein model architecture

In order to improve the runtime complexity of protein structure generation, we use a sparse attention architecture based on a standard transformer with prenormalization and GeGLU update [36, 37]. A schematic overview of this architecture is shown in 1 A. Each block in our model takes as input a set of amino acid features `local`_*i*_ and position features *x*_*i*_. These are fed into a sparse version of invariant point attention (IPA) [21]. Instead of computing the full attention matrix and pair features, we first construct set of neighbours for each amino acid. We extract nearest neighbours based on residue index, proximity in euclidean space as well as additional random neighbours (Fig. 1 A, Sec. 19). Each amino acid only computes pair features and attention weights for its set of neighbours. This procedure reduces attention complexity from *O*(*N* ^2^) to *O*(*N* · *K*), where *N* is the number of amino acids and *K* is the number of neighbours. In contrast to other protein generative models [1, 2, 20] our model does not use persistent pair features with pair attention or triangle multiplication [21], which would increase complexity to *O*(*N* ^3^). We also do not use explicit amino acid frame features [21] that are updated in each block. Instead, our models directly update atom positions and recompute frame information when required to ensure equivariance (Sec. IV B).

**FIG. 1.**
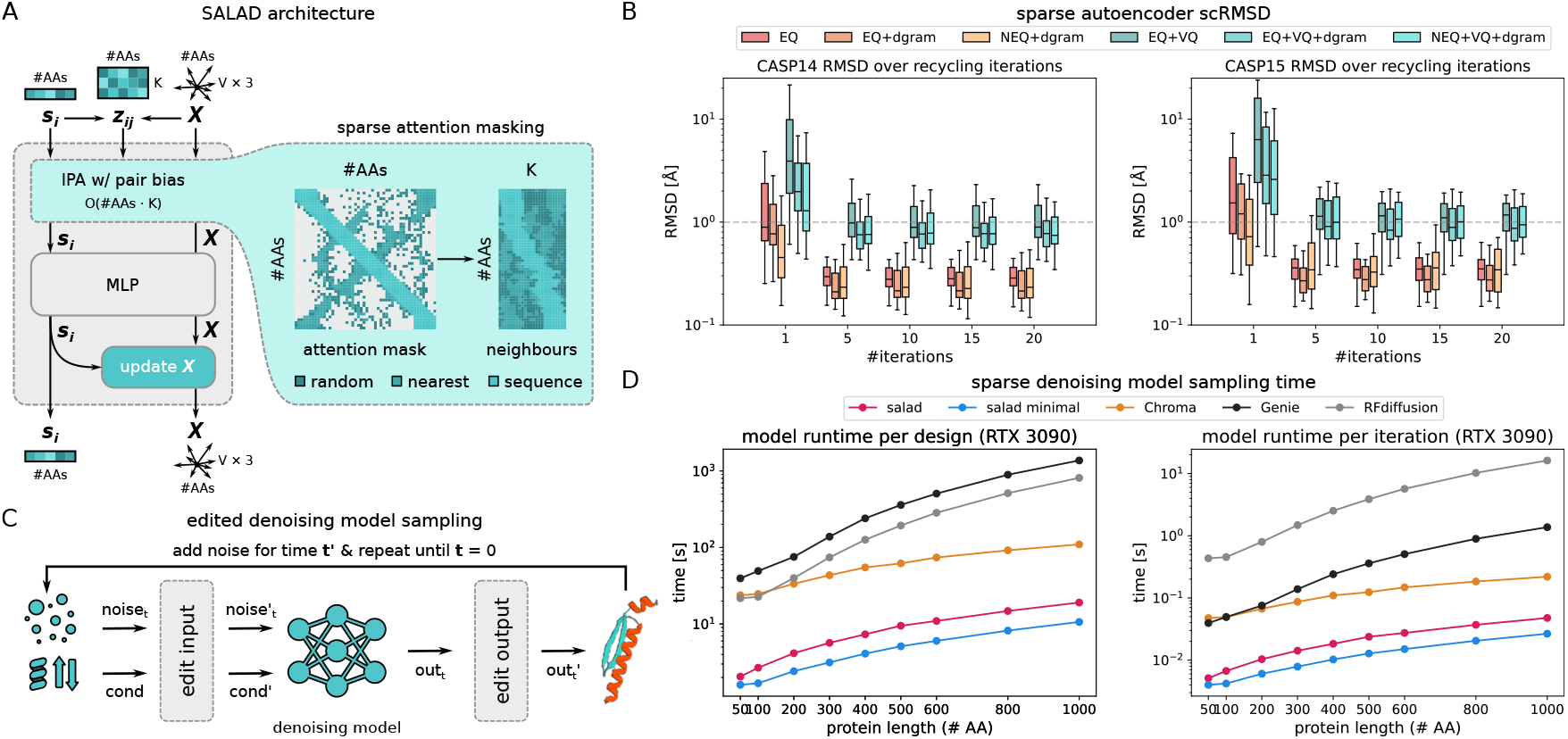
Sparse model architecture and capabilities. (A) Schematic of our sparse protein model architecture. (B) Schematic of the sampling process of a denoising generative model with input and output editing. (C) Sparse model performance on an autoencoding task. Boxplots of scRMSD between ground truth and decoded structures for multiple sparse architectures: equivariant and non-equivariant, using predicted distograms for neighbour selection and selecting neighbours based on current predicted position only. Measures of reconstruction performance are shown per number of model recycling iterations. The dotted line indicates the threshold of 1 Å for reconstruction at atomic precision. The box center line indicates the median, the boundaries indicate the 1st and 3rd quartiles and whiskers show the 1st or 3rd quartile + 1.5× the interquartile range. (D) Average runtimes on a single RTX 3090 GPU by number of amino acids of sparse diffusion models (full pair features and minimal pair features) compared to RFdiffusion, Genie2 and Chroma. Runtimes are reported per designed structure, using the default number of denoising iterations for each model (salad: 400, Chroma: 500, Genie2: 1000, RFdiffusion: 50) as well as the time per iteration.

As a first step to see if our sparse attention architecture can model protein structures, we trained a family of models as autoencoders on proteins in the PDB [38] (Fig. 1 B). We encoded protein structures using a single sparse transformer layer with 32 nearest neighbours per amino acid. We then optionally applied vector quantization (+VQ) to the resulting latent representation [39] and decoded it using a 6-layer sparse transformer with recycling [21]. We tested both equivariant (EQ) and non-equivariant (NEQ) sparse transformers with using different neighbour-selection schemes, either using only euclidean distances, or per-layer predicted amino acid distograms (+dgram) (Section IV D). Evaluating these models on the CASP14 and CASP15 monomer test sets [40, 41] resulted in all models reaching < 1 Å reconstruction accuracy after less than 10 recycling iterations. This indicates that our sparse attention architecture is expressive enough to model protein structures. While there is a large difference in model performance between different architectures at 1 model iteration, this difference decreases with the number of model iterations. As a consequence we decided to move forward with using a simple equivariant architecture without per-layer distogram neighbours for the rest of this work.

### B. Edited denoising protein models

After ensuring that our sparse models are suitable for reconstructing protein structures, we modified our architecture for generating protein backbones. Our models operate on protein structures containing the backbone atoms (N, CA, C, O), an idealized beta carbon (CB) and additional learned pseudo-atoms. We trained our models to denoise noisy structures *x*_*t*_ ∼ *p*(*x*_*t*_|*x*_0_) and to recover the original structure *x*_0_, resulting in a diffusion model loss 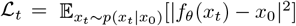 [29]. In addition to recovering *x*_0_, we introduced auxiliary losses to also predict an amino acid sequence and side-chain atom positions. As our models generate all-atom structures, we refer to them as **s**parse **al**l-**a**tom **d**enoising (**salad**) models throughout this work.

At inference we can use our models to generate protein backbones by progressively denoising a pure noise structure *x*_1_. Given a noisy structure *x*_*t*_, we can use the model to predict an estimate *f*_*θ*_(*x*_*t*_) of the denoised structure *x*_0_. Reapplying noise at a lower diffusion time *t*^′^ results in a structure 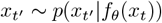 (Fig. 1 C). Repeating this process eventually results in a generated structure *x*. To enforce structural properties of generated backbones directly in the denoising process, we introduce editing functions `edit_input`, `edit_output` which modify the input and output of the denoising model (Fig. 1 C). This results in a generative process:

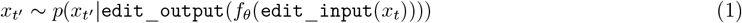

Designing suitable editing functions allows us to adapt our models to various tasks from motif scaffolding (Sec. II E to multi-state protein design (Sec. II G) without having to re-train our models.

As we are using sparse models for the sake of runtime efficiency, we compare the runtime performance of our models to state-of-the-art protein diffusion models (RFdiffusion, Genie2, Chroma). To see how far we can push model runtime, we tested both a full and light-weight version of our model (Sec. 5). For each model, we generated 10 protein backbones per protein length (50-1000 amino acids) on a single nvidia RTX 3090 GPU (Fig. 1 D). salad outperformed all other models in terms of both time per design at the default number of diffusion steps and time per model iteration. In comparison to the fastest non-salad model (Chroma) our models reached up to an order of magnitude of speedup, while outperforming RFdiffusion by up to 2 orders of magnitude on large proteins. Indeed, generating a 1000 amino acid protein structure using salad on a single nvidia RTX 3090 GPU takes only 19 seconds on average, while RFdiffusion takes over 10 minutes. This suggests that we have indeed reached our primary goal of implementing a runtime-efficient protein generative model.

### C. Sparse models generate diverse and designable protein structures

While a flexible sampler and good runtime performance are important properties of our models, we need to assess model performance in terms of the quality of the generated backbones. To compare salad model performance to state-of-the-art diffusion models and hallucination-based approaches, we generated 200 backbones each for proteins of size 50 to 1,000 amino acids (50, 100-600 in increments of 100 amino acids, 800 and 1,000). For each backbone, we designed 8 sequences using ProteinMPNN and predicted their structures with ESMfold. Following current best practices [2, 19] we computed **designability** as the percentage of structures reaching an RMSD between design and predicted structure (scRMSD) < 2 Å and pLDDT > 70 for the best designed sequence (Fig. 2 A). We assessed the impact of different noise distributions on protein structure generation by comparing model performance with both variance preserving (VP) and variance expanding (VE) noise with different standard deviation (80 Å and 100 Å). In addition, we include models trained with protein length-dependent variance VP noise (VP-scaled), as the variance of atom positions in protein backbones varies with the number of amino acids (Sup. Fig. 4 A). We compare the results of our models (VP, VP-scaled, VE) with results previously reported in [2] for RFdiffusion and Genie2 (both models use VP diffusion), as well as results from RSO – the state-of-the-art hallucination-based method for protein design [19] – using the same evaluation approach for all methods.

**FIG. 2.**
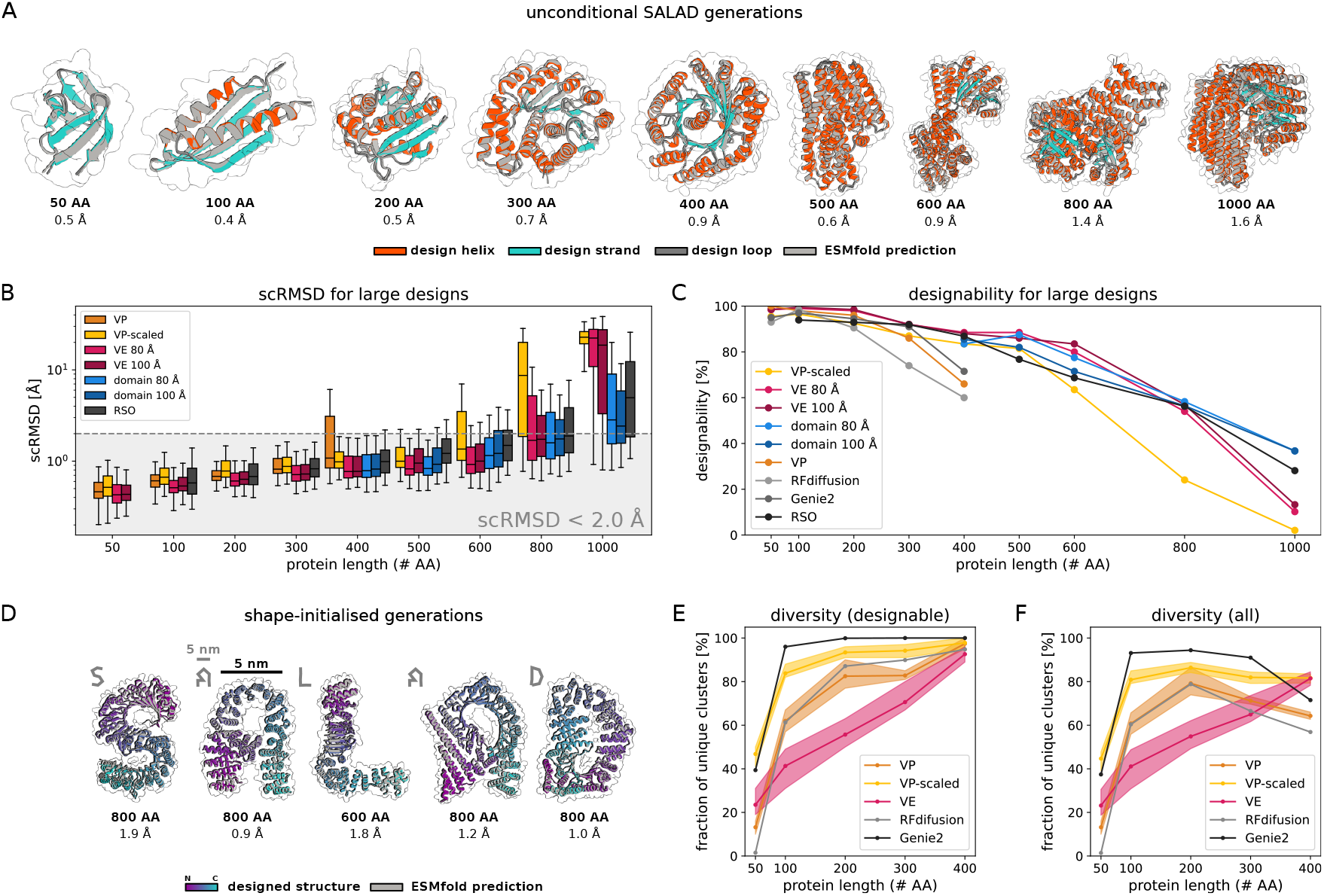
Unconditional structure generation with SALAD models. (A) Example unconditional generations using SALAD, ranging from 50 to 1000 amino acids (coloured by secondary structure; loop: grey, helix: red, strand: blue) and their ESMfold predicted structures (light grey). The scRMSD between generation and ESMfold prediction is listed underneath each structure. (B) scRMSDs using ESMfold for designed structures from 50 to 1000 amino acids. scRMSD distributions for RSO [19] are shown in grey for comparison. The region of successful designs with scRMSD < 2.0 Å is marked in light grey. The box center line indicates the median scRMSD, the boundaries indicate the 1st and 3rd quartiles and whiskers show the 1st or 3rd quartile + 1.5× the interquartile range. (C) Designability of generated structures by protein length from 50 to 1000 amino acids. Values for RSO are shown as reported in [19]. (D) Generations (coloured by residue index) and ESMfold predictions (light grey) for large protein structures generated using VE denoising starting from letter-shaped noise. Each structure is reported with its scRMSD between generation and ESMfold prediction. The grey letter shape in the upper left corresponds to the shape of the noise the proteins were generated from. The grey scale bar corresponds to a distance of 5 nm in each shape. The black scale bar corresponds to the same distance in the depicted protein structures. (E, F) Model diversity computed as the fraction of designable clusters over designable structures (E), or all generated structures (F). Clusters are generated using single-linkage clustering by pairwise TM score with a cutoff of TM-score >= 0.6.

Our models are able to generate designable backbones for a variety of protein lengths from 50 to 1,000 amino acids (Fig. 2 A; Extended Data Fig. 2). Generated structures show low scRMSD, high scTM/pLDDT and diverse secondary structures that include both all-helix and all-strand topologies (Fig. 2 A, B; Extended Data Fig. 3; Extended Data Fig. 5 A, B). In the range of 50 to 400 amino acids, our VP model reaches comparable designability to previous VP models (Genie2, RFdiffusion), outperforming RFdiffusion and slightly underpeforming compared to Genie2, which was trained on a much larger dataset [2] (Fig. 2 C). At 400 amino acids, all VP models show a sharp increase in scRMSD accompanied by a decrease in designability (Fig. 2 B, C). We suspect this decrease in designability is caused by VP diffusion models generating highly compact backbones (Extended Data Fig. 4 A). We find that such backbones require a high fraction of glycine and alanine residues to avoid clashes (Extended Data Fig. 4 B, C), which might decrease designability. A likely cause of this is the fixed variance of the VP diffusion process, which requires the model to reduce amino acid distances at small protein sizes, but increase amino acid distances at large protein sizes (Fig. 4 D). If the model trains predominantly on small proteins, this discrepancy might result in the observed compact backbones for larger proteins at inference. This issue with protein diffusion models is anecdotally known to the protein design community [42].

In contrast, VP-scaled and VE models do not experience increases in scRMSD at the 400 amino acid threshold. VP-scaled models maintain median scRMSD < 2 Å for proteins of up to 600 amino acids, whereas VE models maintain this value for proteins up to 800 amino acids in length (Fig. 2 B). This is mirrored by designability, where both VP-scaled and VE models outperform all VP models at protein lengths above 300 amino acids. However, neither VP-scaled nor VE models can maintain high designability for generated backbones of length 1,000 where both types of models drop below 20 %. We hypothesized that this decrease in designability is due to the models being unable to properly model the global structure of large proteins. As large proteins generally consist of multiple domains [43], we tested if VE models initialized from domain-shaped noise (Sec. IV K) would result in lower scRMSD and greater designability for large proteins. Instead of using normal distributed noise centered on the coordinate origin, we first sample a set of domain centers and then add normal distributed noise (with standard deviation 80 Å or 100 Å) for 200 amino acids to each of these centers (Sec. IV K). At every subsequent denoising step, we use standard VE noise. Using this noise initialization leads to decreased scRMSD and increased designability for large proteins, reaching a designability of up to 36.7 % for 1,000 amino acid proteins (Fig. 2 C). This way, domain-shaped noise matches or improves on the designability of RSO [19], the current state-of-the-art approach to large protein design (Fig. 2 B, C).

As we can use domain-shaped noise to generate large proteins with VE models, we investigated if we could control the shape of generated backbones using noise with a specified shape. By sampling the initial noise centered on letter shapes, we were able to generate designable structures spelling out the name of our framework (Fig. 2, D). In contrast to previous work on shape-conditioned protein design using the Chroma model [20], our approach does not require an additional shape conditioner and results in designs with low scRMSD and high pLDDT (Fig. 2 D). In terms of standard designability criteria using ESMfold (scRMSD < 2 Å, pLDDT > 70), 55 % of letters generated by our models are designable, whereas up to 92.5 % of letters are refoldable according to the criteria used for Chroma (scTM > 0.7) [20] (Extended Data Fig. 1). This indicates that our models can be used to generate designable backbones even on challenging out-of-distribution design tasks.

In addition to designability, we measure the **diversity** of protein backbones generated by our models. Lin *et al*. [2] have previously defined **diversity** of generated backbones by single-linkage clustering designable backbones by TM score with a cutoff of TM > 0.6. Diversity is then computed as the fraction of designable clusters in all generated backbones 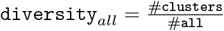 [2]. This diversity measure implicitly includes backbone designability, as a lower number of designable backbones results in a lower number of clusters. A method trading off designability for increased diversity would therefore result in a low `diversity`_*all*_ score. To disentangle diversity and designability, we decompose this diversity score as

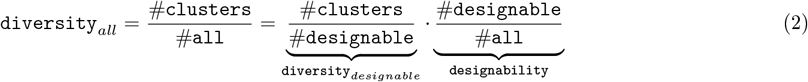

When computing `diversity`_*designable*_, only the diversity of designable structures is taken into account and diversity is not deflated by low designability. We argue that this is a more meaningful measure of diversity as only designable structures are used for protein design in the end.

We compare our models with RFdiffusion [1] and Genie 2 [2]. For proteins of length 50 to 400 amino acids we take random samples of 100 generated structures and compute both `diversity`_*all*_ and `diversity`_*designable*_ for each sample. To quantify the spread of diversity across samples, we show both the median as well as the minimum and maximum diversity over 10 samples (Fig. 2 E, F). Our VP model achieves similar diversity to RFdiffusion for both `diversity`_*designable*_ and `diversity`_*all*_, while the VP-scaled model outperforms RFdiffusion on both metrics and approaches the diversity of Genie 2 outperforming it in terms of `diversity`_*all*_ for 400 amino acid proteins. Our VE model shows reduced diversity at small protein sizes but shows comparable `diversity`_*designable*_ to RFdiffusion on 400 amino acid proteins and outperforms both Genie2 and RFdiffusion in terms of `diversity`_*all*_ for this protein length. This indicates that designable 400 amino acids structures generated by our non-VP models are comparably diverse to those generated using Genie2. Their increased `diversity`_*all*_ can be attributed to their improved designability (Fig. 2 C). We therefore argue that `diversity`_*designable*_ is a more meaningful measure of diversity as it is not inflated by changes in designability. While our models sightly underperform Genie2 in terms of designability, we note that Genie2 was trained on AlphaFold DB [44] - a larger and more diverse dataset.

### D. Random secondary structure conditioning maximizes diversity

As our models can be conditioned to generate proteins with a given secondary structure, we investigated if conditioning models with random secondary structures could increase the diversity of generated backbones. We sampled random three-state secondary structure strings (helix, strand, loop) by selecting a random percentage of helices and strands, constructing secondary structure elements of random lengths that add up to the selected percentages and randomly arranging them into a secondary structure string (Fig. 3 A). We then used our denoising models to produce backbones for each random secondary structure. Computing `diversity`_*designable*_ for backbones of length 50 to 400 amino acids generated this way resulted in our models surpassing RFdiffusion at all sizes and matching or outperforming the designability of Genie2 while being trained on a much smaller dataset [2] (Fig. 3 B). Notably, random secondary structure conditioning resulted in an increased diversity for small proteins and saturated the diversity metric on proteins of length 200 or larger. However, increasing diversity this way resulted in decreased designability across all protein lengths and models (Extended Data Fig. 5 C).

**FIG. 3.**
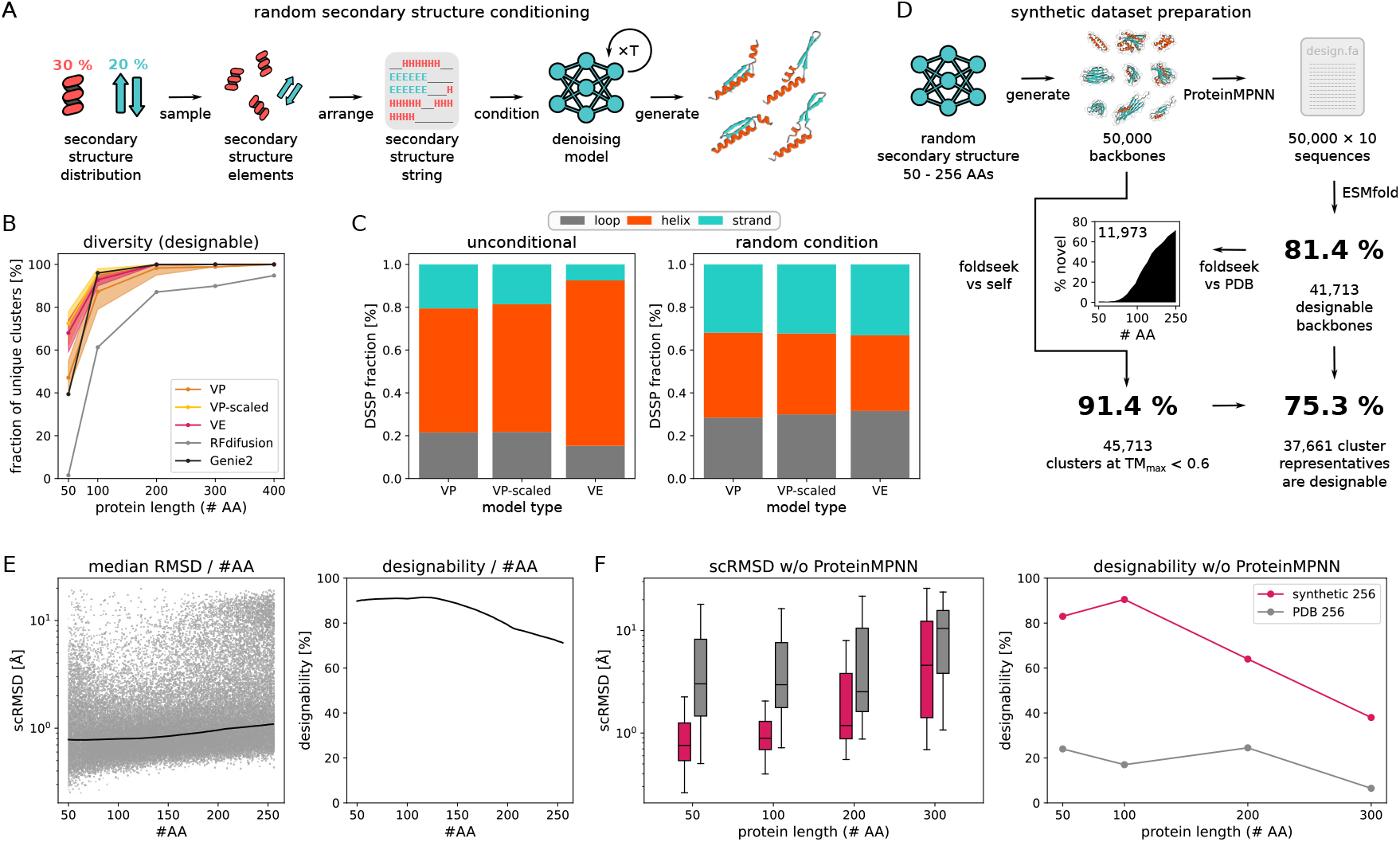
Random secondary structure conditioning maximises diversity at the cost of designability. (A) Schematic of our random secondary structure sampling procedure. (B) Diversity of designable structures generated using random secondary structure conditioning for sizes 50 to 400 amino acids, compared to the diversity of designable structures generated using RFdiffusion and Genie2. (C) Secondary structure distribution for our models using no conditioning (left) or random secondary structure conditioning (right). (D) Overview of diverse synthetic dataset generation using random secondary structure conditioning. (E) Left: Scatter plot of ESMfold scRMSD for all 50,000 generated structures. The line indicates the median scRMSD within a length-window of 100 amino acids. Right: designability of generated structures in the synthetic dataset computed for a length-window of 100 amino acids. (F) Single-shot performance of diffusion models trained on the synthetic dataset (synthetic 256) compared to the subset of proteins of length < 256 amino acids in PDB (PDB 256). Left: Boxplot of RMSD between generated structures and ESMfold predictions for the argmax sequence prediction for models trained on the synthetic dataset and PDB. The center line indicates the median scRMSD, box boundaries the 1st and 3rd quartiles and whiskers 1.5× the inter-quartile range from the box. Right: Designability of argmax sequence predictions for models trained on synthetic data and PDB.

**FIG. 4.**
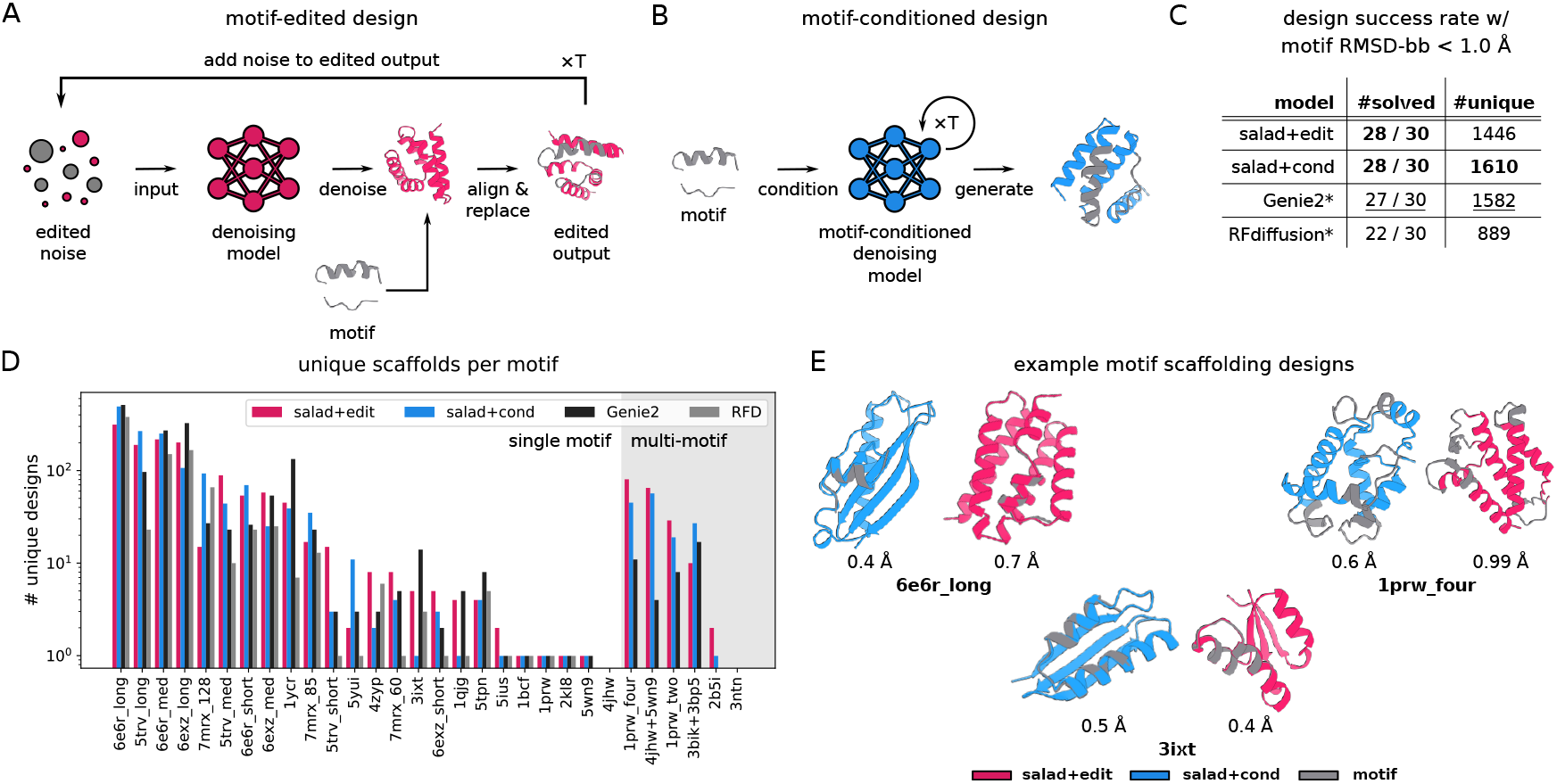
Motif scaffolding. (A) Schematic of the output-editing procedure for motif scaffolding. (B) Schematic of model conditioning for motif scaffolding. (C) Table of unique successful designs on the motif scaffolding benchmark established by Lin *et al*. 2024 [2] using output editing and model conditioning. Results for RFdiffusion and Genie2 are show as reported by Lin et al. 2024 [2]. (D) Bar plot of the number of unique successful designs (as measured by single-linkage clustering at TM score < 0.6) for salad models with editing and conditioning, compared to results reported in Lin et al. 2024. All evaluations were performed with the same settings as Lin et al. 2024 [2]. (E) Example structures of scaffolded motifs using output editing and model conditioning. All displayed structures are ESMfold predictions of designed sequences with the motifs marked in grey. Motif RMSD is reported below each structure.

In addition to greatly increasing clustering-based diversity, this approach also equalized secondary structure content biases inherent to our models (Fig. 3 C; Extended Data Fig. 5 A, B, D). While all of our models showed a preference for alpha-helices for unconditional generation, conditioning resulted in a uniform distribution of secondary structure content. Quantifying the diversity in secondary structure content of designs showed that conditioning increased the entropy of the secondary structure distribution relative to the non-conditioned baseline (Extended Data Fig. 5 E, F). This indicates that conditioned designs are more diverse both in terms of shape and secondary structure content.

To test the limits of random secondary structure conditioning for generating diverse protein structures, we generated a synthetic dataset of 50,000 backbones with size between 50 and 256 amino acids (Fig. 3 D; Section IV M). We designed 10 sequences per backbone with ProteinMPNN, predicted their structures with ESMfold and quantified designability, diversity as well as novelty with respect to the PDB. Of the 50,000 backbones, 81.4 % were designable. Across protein sizes, designs showed low median scRMSD and high overall designability (Fig. 3 E). To quantify diversity, we clustered all backbones using foldseek with TMalign alignment (TM-score threshold 0.6) and a minimum coverage of 90 % of the sequence to only cluster structures of similar sizes [45]. This yielded 45,713 clusters corresponding to 91.4 % of the dataset. Of these cluster representatives, 75.3 % were designable, resulting in a dataset of 37,661 diverse and designable structures. Using foldseek to search the PDB for matches for all designable structures in the dataset resulted in 11,973 structures without a single match at TM-score > 0.5. Notably, most matches were concentrated in short backbones, with the majority of backbones with 200 or more amino acids had no matches in the PDB. This indicates that generating structures with random secondary structure can explore parts of protein fold space far outside the training set and result in “dark matter” folds outside the PDB.

Previous work on protein generative models [46] reported that training on synthetic data with ProteinMPNN-designed sequence could improve model performance. To check if a synthetic dataset generated this way could be used to potentially train improved protein generative models, we compared the performance of two `salad` models trained on proteins of size 50 to 256 amino acids. We trained one model on a subset of PDB with chains of length between 50 and 256. The other was trained on designable structures and sequences in our synthetic dataset. We generate 200 backbones for protein sizes between 50 and 300 amino acids. As our models learn to predict a sequence as an auxiliary task during training (Section IV C), we generated a single sequence per backbone. We predicted the structure of each sequence using ESMfold [22] to assess design success. The model trained on PDB resulted in high median scRMSD (> 2 Å) and low designability (< 20 %) across all tested protein sizes (Fig. 3 F). In contrast, the model trained on our synthetic dataset showed low median scRMSD and high designability for in-distribution tasks, with performance deteriorating for proteins of size 300 amino acids, which the model was not trained on (Fig. 3 F). Directly generating successful backbone-sequence pairs circumvents the sequence design step in the protein design pipeline, reducing the number of tested sequences and AlphaFold or ESMfold evaluations for design filtering from 8 to 1. This greatly decreases the runtime of the protein design pipeline.

### E. Structure-editing for motif scaffolding

While unconditional backbone generation can give an indication about the general performance of a protein generative model, it is rather removed from realistic applications of protein generative models. Motif-scaffolding provides a more realistic benchmark task. Models have to generate backbones that accommodate one or more functional motifs from natural proteins [1]. This has immediate applications to enzyme design (scaffolding theozymes) [15, 16], synthetic vaccine design [7] and the design of natural protein mimics [47].

We compare the performance of `salad` models against the state-of-the-art protein diffusion models Genie2 and RFdiffusion on a standardized motif-scaffolding benchmark. The benchmark, introduced in [1] includes 24 single-motif tasks of varying difficulties and was extended in [2] to contain an additional 6 tasks where the models have to scaffold more than one motif in a single backbone (multi-motif scaffolding). For direct comparison to Genie2 and RFdiffusion, which are both VP models, we only use VP models in this benchmark. As our models are not trained for multi-motif scaffolding by default, we approach this problem in two different ways. First, we use our structure-editing approach (Section II B) to edit the denoised structure by aligning the motif backbone and replacing the output coordinates by the motif’s coordinates (Fig. 4 A). This ensures that the motif is present in the final generated backbone, even if the model is not conditioned on the motif’s structure (Section IV P). We call this configuration `salad+edit`. Second, we train a separate multi-motif conditioned model which we will refer to as `salad+cond` (Fig. 4 B; Section IV O).

In the following we compare the results for our method with the results for RFdiffusion and Genie2 reported in [2]. To directly compare with these, we closely followed the same evaluation strategy. For each approach, we generated 1,000 backbones per scaffolding problem, designed 8 sequences with ProteinMPNN and assessed designability with ESMfold (Section IV Q). We further filtered designable structures by their motif RMSD computed over all backbone atoms (N, CA, C, O). Structures were deemed successful if they reached motif RMSD < 1 Å. All successful structures were then clustered using TMalign at a TM-score cutoff of 0.6 to identify unique scaffolds for each problem. Evaluating success using CA-based motif RMSD showed little to no impact on both the number of successful and unique designs for `salad+cond` (Fig. 7 A, B). In contrast, `salad+edit` showed large variability in success rates for some motifs (Fig. 7 A, B). This indicates that the lack of explicit motif conditioning may result in the model changing motif orientation in the denoising step.

We found that both `salad+edit` and `salad+cond` solved 23/24 single-motif as well as 5/6 multi-motif design tasks (Fig. 4 C, D). Both `salad+edit` and `salad+cond` generated diverse backbones with low motif RMSD (Fig. 4 E). Only the motifs for 4jhw and 3ntn remained non-designable, consistent with Genie2 [2]. However, compared to Genie2 we were able to solve one additional multi-motif scaffolding task with 2b5i. While RFdiffusion cannot be straight-forwardly applied to multi-motif scaffolding, our approaches still outperformed it on single-motif scaffolding, solving one additional problem. Overall, `salad+cond` generated 1,610 (`salad+edit` 1,446) unique scaffolds, slightly outperforming Genie2 and dwarfing RFdiffusion’s 889 scaffolds (Fig. 4 C).

Thus, our models outperform RFdiffusion across all criteria, while matching Genie2 in terms of the total number of unique scaffolds and solving one additional problem with 2b5i. While a direct comparison of the number of unique backbones per scaffolding problem (Extended Data Fig. 7 C) shows that there is currently no best model across all tasks, our models result in equal or more scaffolds for the majority of design tasks (21 / 24 vs RFdiffusion and 19 / 30 vs Genie2 for output editing; 20 / 24 and 20 / 30 for conditioning) (Fig. 7 C). This indicates that both our approaches are competitive with the state-of-the-art for single and multi-motif scaffolding.

### F. Structure-editing for repeat protein design

As a second application to demonstrate the flexibility of our models combined with output editing, we set out to generate repeat proteins. Similar to the approach used in previous work [1], we can generate point symmetric repeat proteins by symmetrizing the inputs of our models according to the action of a point group (Fig. 5 A). As our model has amino acids attend to random neighbours (Section 19), we also symmetrize the output of our models. All repeat subunits are aligned using the action of the point symmetry group and averaged to produce a representative repeat unit. This can then be placed at a specified radius from the symmetry axis to control the radius of the generated symmetric repeats. Replicating the representative structure then gives us a symmetrized output structure (Section IV R). Applying this procedure resulted in designable backbones with low scRMSD for various cyclic symmetry groups (Fig. 5 B).

**FIG. 5.**
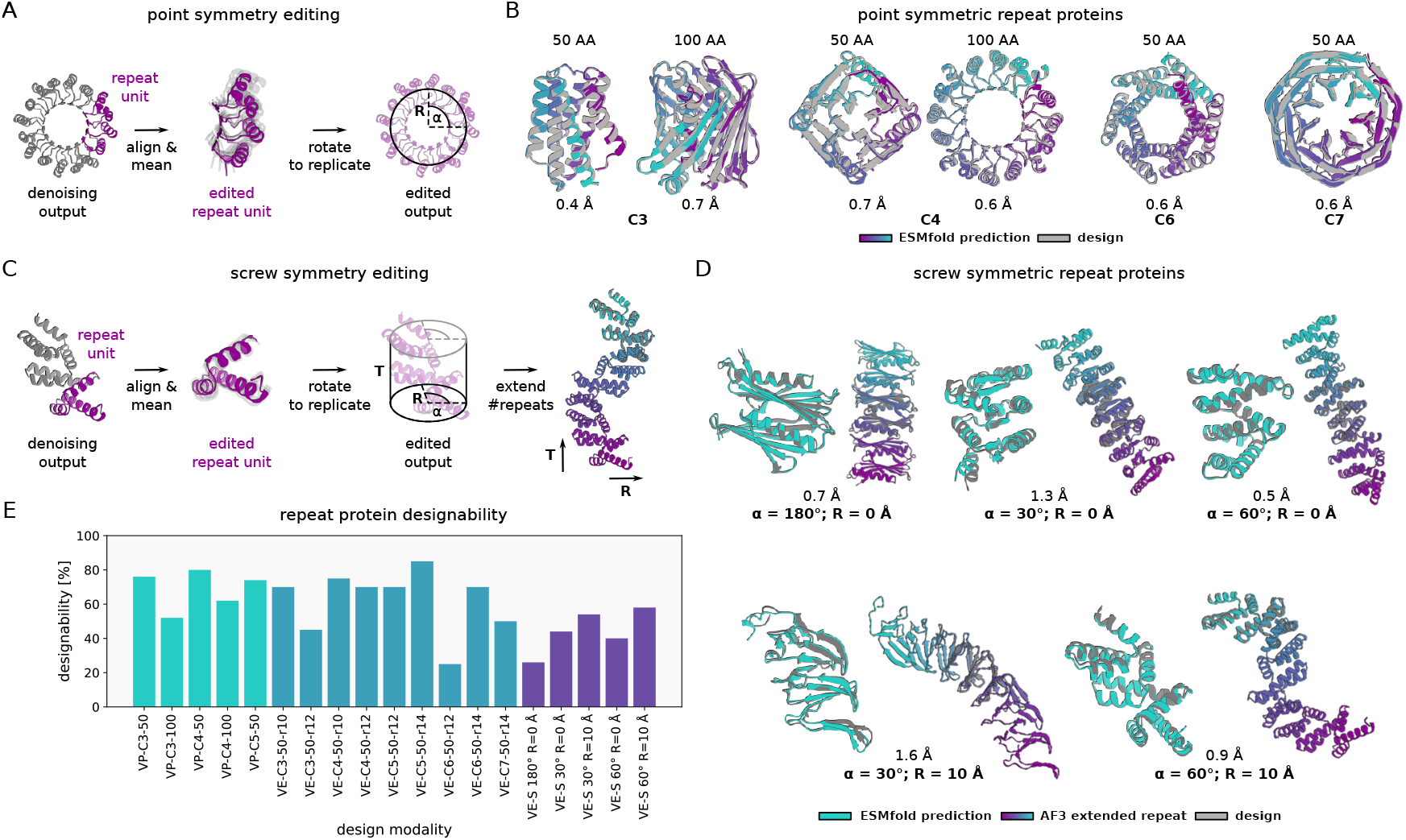
Edited denoising for symmetric and repeat proteins. (A) Schematic of the output editing procedure for point symmetric repeat protein design. (B) Example generated point symmetric repeat proteins for different subunit sizes (50, 100 amino acids) and cyclic groups (C3, C4, C6, C7). The idealized symmetric design is shown in grey, an ESMfold prediction for each design coloured by residue index. scRMSDs are reported for each example structure. (C) Schematic of the output editing procedure for screw-symmetric repeat proteins. (D) Example screw-symmetric designs (grey) for different angles (30° - 180°) and radii (0 Å, 10 Å), with a fixed inter-subunit translation of 12 Å. An ESMfold predicted structure (turquoise) is superimposed onto the designed repeat (left). Additionally, an AF3 [48] predicted structure of the design replicated 3 times (coloured by residue index) is shown together with the scRMSD of the design to that extended repeat protein. (E) Design success-rates for cyclic and screw-symmetric designs using VP and VE models with different symmetry groups, radii and rotation angles.

While both hallucination [5] and diffusion [1] methods have been successful in producing point-symmetric proteins, only sequence-based [13] machine learning methods and Rosetta-based protein design [3, 4], have been successfully applied to design extended repeat proteins that cannot be described by a point group. However, our editing approach is readily extended to arbitrary symmetry groups including screw (helical) symmetries, which cover the class extended repeats described in [3, 4, 13] (Fig. 5 C). By explicitly setting the radius *R*, rotation angle *α* and translation *T* of a screw symmetric repeat (Fig. 5 C), we can generate designable backbones with the specified geometry (Fig. 5 D). Generated backbones are structurally diverse, with topologies ranging from fully alpha-helical to fully beta-sheet. Repeat sequences designed using ProteinMPNN [23] are reliably predicted to take on the designed structure with low scRMSD, even when extending the number of repeats by a factor of 3 (Fig. 5).

Overall success rates for symmetric protein design vary between models (VP or VE) and design tasks (Fig. 5 E). Strikingly, designabilities for both cyclic and screw symmetric designs show a dependency on the specified radius and rotation angle, e.g. C6 symmetry with radius 12 Å (“r12”) and radius 14 Å (“r14”). This is likely due to generated structures becoming highly compact for low radii, which is associated with a loss in designability (Section II C; Extended Data Fig. 4).

### G. Structure-editing for multi-state design

While both motif scaffolding and repeat protein generation demonstrate the applicability of output editing to protein design, these tasks do not showcase the full flexibility of this approach. In both cases, external conditioning information is available in the form of a motif or symmetry group. Additionally, there is sufficient data to train structure generative models conditioned on either of these tasks. In contrast, designing amino acid sequences which can fold into multiple distinct backbone structures (multi-state design) fits none of these criteria: data on natural multi-state proteins is scarce [49] and structure-based generative models are believed to be unsuitable for this task [13]. Therefore we chose to demonstrate that `salad` models can solve a recent multi-state benchmark task introduced in [13] by using output editing to couple the outcome of multiple denoising processes – one for each state (Section IV T).

Following [13], we designed backbones for a protein (the “parent”) that takes on a specified secondary structure when intact and a different secondary structure when split into two “child” proteins. The N and C terminal parts of the parent share their secondary structure with the children, while the central part of the parent should transition from a beta-sheet to an alpha helix when split (Table III). As we cannot directly design the protein on the sequence level, we instead instantiate 3 separate denoising processes, one for each state (Fig. 6 A). Each denoising process is conditioned on the secondary structure string of either parent, child 1 or child 2. At each denoising step, we enforce that the parts with the same secondary structure between parent and child share a similar 3D geometry by aligning and averaging their substructures (Fig. 6 A; Section IV T). Essentially, the denoising processes are coupled by conditioning their structures on each other. This approach results in 3 coupled structures per generation.

**FIG. 6.**
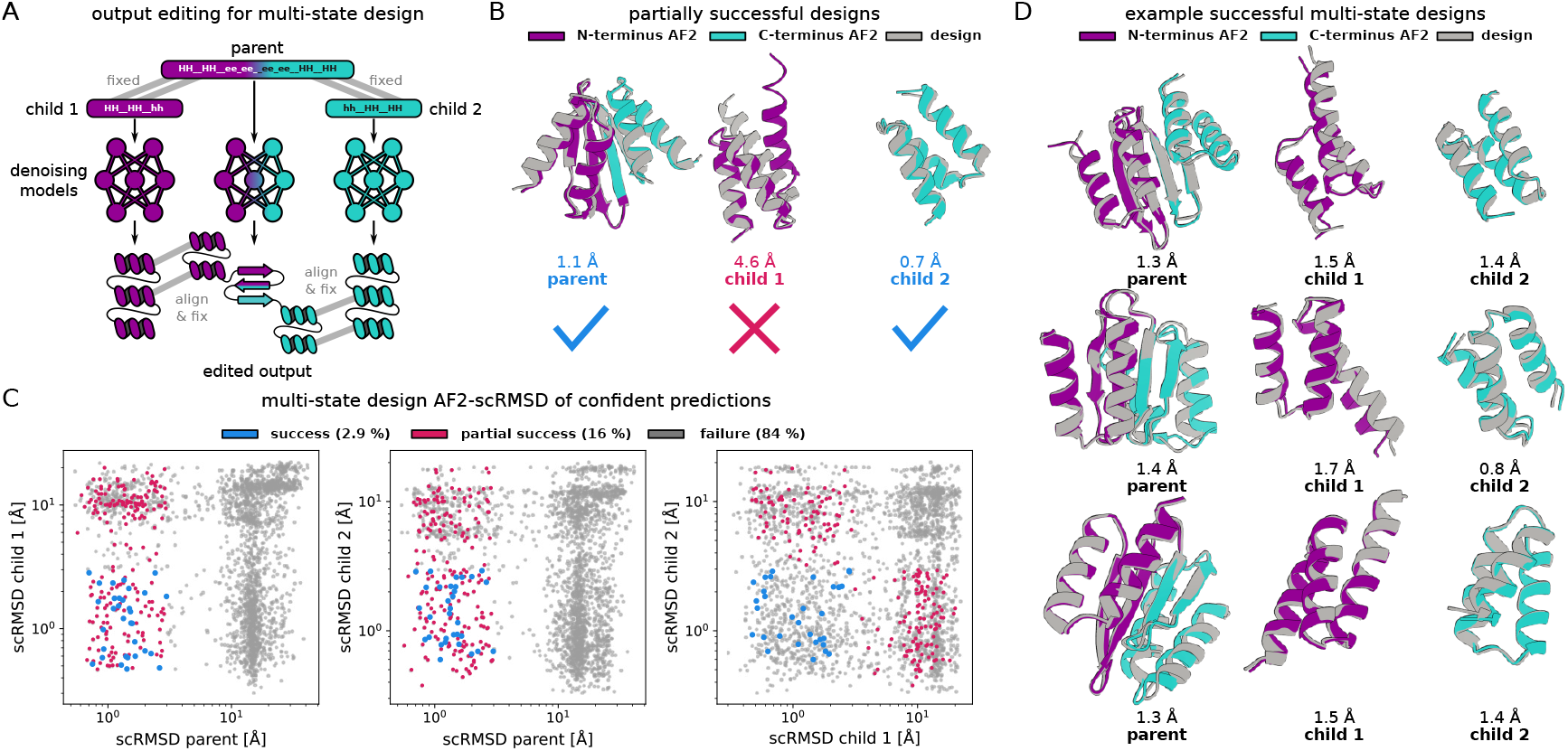
Multi-state protein design. (A) Schematic of the output editing procedure for multi-state protein design. (B) Example partially successful multi-state design. (C) Scatter plots of AF2 scRMSDs for parent and child sequences with pLDDT > 75. Failed sequences are marked in grey, partial successes (parent and at least one child successful) are marked in purple and successful designs are marked in blue (D) Examples of successful multi-state designs. Generated structures for parent and child proteins are shown in grey; predicted structures are coloured by their corresponding child protein. The N-terminal part is coloured purple and the C-terminal part is blue. AF2 scRMSDs are reported for each parent and child design.

We generated 1,000 structure triples, designed their sequences with ProteinMPNN, tying the sequence across parent and child structures [23]. To compare with the benchmark done in [13], we evaluated design success with the same criteria of AlphaFold RMSD < 3.0 Å and pLDDT > 75. 16 % of generated backbones resulted in partially successful designs, where all structures passed the pLDDT threshold, and both the parent and at least one child passed the RMSD threshold (Fig. 6 B). While these designs did not meet all criteria, they still resulted in a sequence predicted to change conformation in the parent compared to the child state. Using the complete criteria, 2.9 % of backbones generated by our approach were successful, compared to the 0.05 % of successes reported for ProteinGenerator (PG) in [13]. While we used ProteinMPNN for sequence design, PG returns one sequence per backbone. However, even evaluating the percentage of success on a sequence level `salad` reached a success rate of 0.32 %, outperforming PG by a factor of 6.

To check whether output editing contributed to the success rate of multi-state design, we generated another 1,000 backbone triples using only secondary structure conditioning. This resulted in per-backbone success rates of 0.4 % and a per-sequences success rate of 0.06 % matching PG (Extended Data Fig. 8). This indicates that structure-editing is indeed behind the increased performance observed for this multi-state design task. Overall, our structure-editing approach generates varied low-RMSD solutions to this multi-state design problem (Fig. 6 D), outperforming previous machine learning approaches. To our best knowledge, this is the first demonstration of multi-state design with a protein backbone denoising model.

## III. DISCUSSION

In this work we present `salad`, a family of efficient sparse denoising models capable of generating designable and diverse protein structures up to a length of 1,000 amino acids. In terms of unconditional protein structure generation, our models outperform RFdiffusion both in terms of designability and diversity for all protein lengths, while closely approaching the diversity of Genie2 which was trained on a larger and more diverse dataset. We bridge this gap in diversity by applying secondary structure conditioning at the cost of designability. This allows us to generate a large, highly-diverse data set of designable structures novel to the PDB. For longer proteins from 400 to 1,000 amino acids, our models decisively outperform both RFdiffusion and Genie2, approaching the designability observed for hallucination-based methods [19]. To our knowledge, `salad` is the first model to bridge this performance gap between protein diffusion models and hallucination-based methods [19] for large protein structures. While achieving high designability and diversity across protein lengths, `salad` models greatly reduce time-per-design due to their efficient sparse architecture.

We expand the capabilities of `salad` by combining it with structure-editing. By editing the output of a `salad` model at each denoising step, we can rapidly prototype generators for protein design tasks unseen during training. We show that this combination can generate designable, low-RMSD backbones for a variety of tasks. We design shape-conditioned proteins, scaffold multiple functional motifs, generate repeat proteins and produce protein sequences predicted to adopt a distinct folds when cleaved. For motif scaffolding, we match or exceed the performance of both RFdiffusion and Genie2. For repeat protein design, we generate screw-symmetric repeats which to our knowledge have not been explored using protein structure diffusion models beyond a single mention in the supplement of Chroma [20]. Instead, such proteins have so far been designed using Rosetta [4] or sequence-based design methods [3, 13]. For multi-state protein design, we reproduced a design task introduced in [13] and achieved a success-rate one order of magnitude higher than the original work. This indicates that `salad` is in fact sufficiently flexible to generate designs even for tasks like multi-state design which are believed to be unfavourable for structure generative models [13].

While `salad` produces acceptable results on computational benchmarks, this work does not contain additional experimental validation. However, the models `salad` is compared to, in particular RFdiffusion [1] and ProteinGenerator include extensive experimental validation. We argue that `salad` matching or exceeding their performance in terms of ESMfold / AlphaFold design success should alleviate concerns about the lack of experimental validation. In particular, `salad` is part of the same pipeline of structure generation, ProteinMPNN sequence design and AF2 / ESMfold selection as RFdiffusion [1]. As `salad` was trained independently from ProteinMPNN and AF2 / ESMfold it is unlikely that it has learned to generate backbones that are adversarial for both. We also emphasize that the focus of this work is on developing more efficient and versatile backbone generators, not to present an all-in-one solution for protein design.

Another limitation of `salad` is that it is currently restricted to a limited training set. `salad` is trained on protein structures in the PDB, with all small molecules, ions, waters and nucleic acids removed. Therefore its uses for enzyme design and small molecule binder design are limited. In particular, more recent versions of RFdiffusion can design proteins in the presence of small molecules [16, 50]. The `salad` architecture is likely capable of handling small molecules with minor modifications which makes this an obvious next step to address in future work. In addition, `salad` struggles to match the diversity of Genie2 [2] without using random secondary structure conditioning. Genie2 was trained on a clustered subset of the AlphaFold database, which greatly exceeds our PDB data set in both size and structural diversity [2, 44, 51]. We believe that this issue can be relatively easily addressed in future work by training `salad` models on AlphaFold DB.

In this work, we largely compare `salad` to RFdiffusion [1] and Genie2 [2]. While many other protein diffusion models exist [31, 52–57], we argue that comparing to these ones in particular is sufficient to establish `salad` as on par with the state-of-the-art, as prior work has shown both Genie2 and RFdiffusion outperform most other protein structure generative models [2]. In particular, the designability of `salad` generations for large proteins as well as its runtime performance give it an edge over comparable models. This leads us to believe that `salad` can fill the niche it was designed for and provide efficient and versatile backbone generation in the first step of the protein design pipeline.

## IV. METHODS

### A. Protein structure denoising models

Denoising protein structure models are trained to reconstruct a noise-free structure *x* from a noisy input structure. To train our models, we sample random time points *t* ∈ [0, 1], where *t* = 0 corresponds to a noise-free structure and *t* = 1 corresponds to pure noise. Depending on the noise schedule we then convert *t* into a noise-scale *σ*_*t*_. We use 3 different noise schedules: variance preserving noise with a cosine schedule [58] and constant standard deviation noise (VP); VP noise with a protein size-dependent standard deviation (VP-scaled); variance expanding noise (VE).

For VP noise, a noisy structure *x*_*t*_ is then generated by sampling

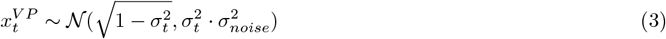

where *σ*_*noise*_ = 10Å is the standard deviation of the noise at time *t* = 1. For VP-scaled noise we instead set the standard deviation *σ*_*noise*_ equal to the standard deviation of CA positions in input structure *x*:

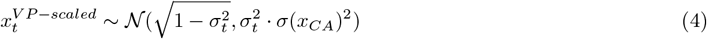

The fully noised structure 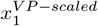 will therefore have the same standard deviation as the alpha carbons starting structure *x*_*CA*_. Finally, for VE noise, instead of sampling a diffusion time *t*, we directly sample a noise-scale from a log-normal distribution, following [59]

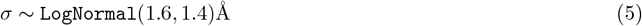

and sample noisy structures according to:

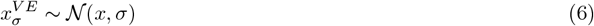

The models are then trained to reconstruct *x* from *x*_*t*_ by minimizing 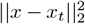 and additional auxiliary losses (Section IV C).

Our models are trained using self-conditioning. At each training step we sample two noisy structures *x*_*t*_, 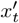. With probability 50 % we predict *x*^′^ from 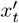 without self-conditioning. We then predict *x* from *x*_*t*_ using *x*^′^ as an additional input. At inference, the model is passed the current noised structure *x*_*t*_ as well as its previous prediction *x*_*prev*_ to predict *x*.

In addition to self-conditioning, our models can also be conditioned on amino acid sequence, partial structure information, three-state secondary structure (helix, strand, loop), block contacts between secondary structure elements, inter-chain contact information and hotspot residues interacting with other protein chains [1]. During training, conditioning information is provided at random for 50 % of training examples. Each conditioning modality is further randomly masked for a random fraction between 20 and 100 % of residues. To determine inter-residue and inter-chain contacts for conditioning, we compute pairwise CA distances between amino acids. Amino acids are considered in contact, if their CA distance is < 8 Å. Two chains and secondary structure elements are considered in contact if at least one pair of residues is in contact. Residues are considered hotspot residues if they are in contact with at least one residue in another chain. Partial structure information is presented to the model as a matrix of inter-residue CB distances together with a mask of amino acid pairs with valid conditioning information. Partial structure information is masked out for between 20 % and 80 % of residues in any training example.

To sample from a trained denoising model, we initialize a backbone *x* with all atom positions set to 0. We then partition the interval [0, 1] into *N* equally-spaced time-steps (*t*_0_, …, *t*_*N*_) with *t*_*N*_ = 1. For each time-step *t* starting with *t*_*N*_, we apply noise with a chosen noise schedule to *x*, resulting in a noised structure *x*_*t*_. This noisy structure is denoised by the model resulting in a new structure *x*. Repeating this process gradually reduces the noise level and results in a denoised protein backbone. Our approach differs from the denoising processes described in the literature: protein structure denoising diffusion models generally sample structures *x*_*t*_ according to a distribution *q*(*x*_*t*_|*x*_*t*+*s*_, *x*) which depends both on the denoised structure *x* and an earlier noisy structure *x*_*t*+*s*_ at diffusion time *t* + *s* [1, 2, 29]. Instead, our approach samples from *q*(*x*_*t*_|*x* = *f*_*θ*_(*x*_*t*+*s*_)), removing any direct dependency on the previous noise *x*_*t*_ and only depends on it through the model *f*_*θ*_. This approach has been previously reported for categorical text diffusion models [60] and more recently amino acid sequence diffusion models [13].

We chose this approach not because of its success in sequence diffusion models, but to enable arbitrary modification of denoised structures *x* without having to take into account *x*_*t*_. To use our models for protein generation tasks they were not trained for, we want to allow arbitrary editing of the denoised structure – for symmetrization, to introduce structural motifs for scaffolding and to couple multiple denoising processes for multi-state design. This necessitates translating, rotating and replacing parts of the denoised structure (see Sections IV P, IV R, IV T). Changing the denoised structure *x* this way without also adjusting *x*_*t*+*s*_ in a compatible way could result in failure to generate valid protein structures.

Algorithm 1 shows the generative process for a model involving conditioning information *c*, self-conditioning and structure-editing.

#### Algorithm 1

Denoising model inference

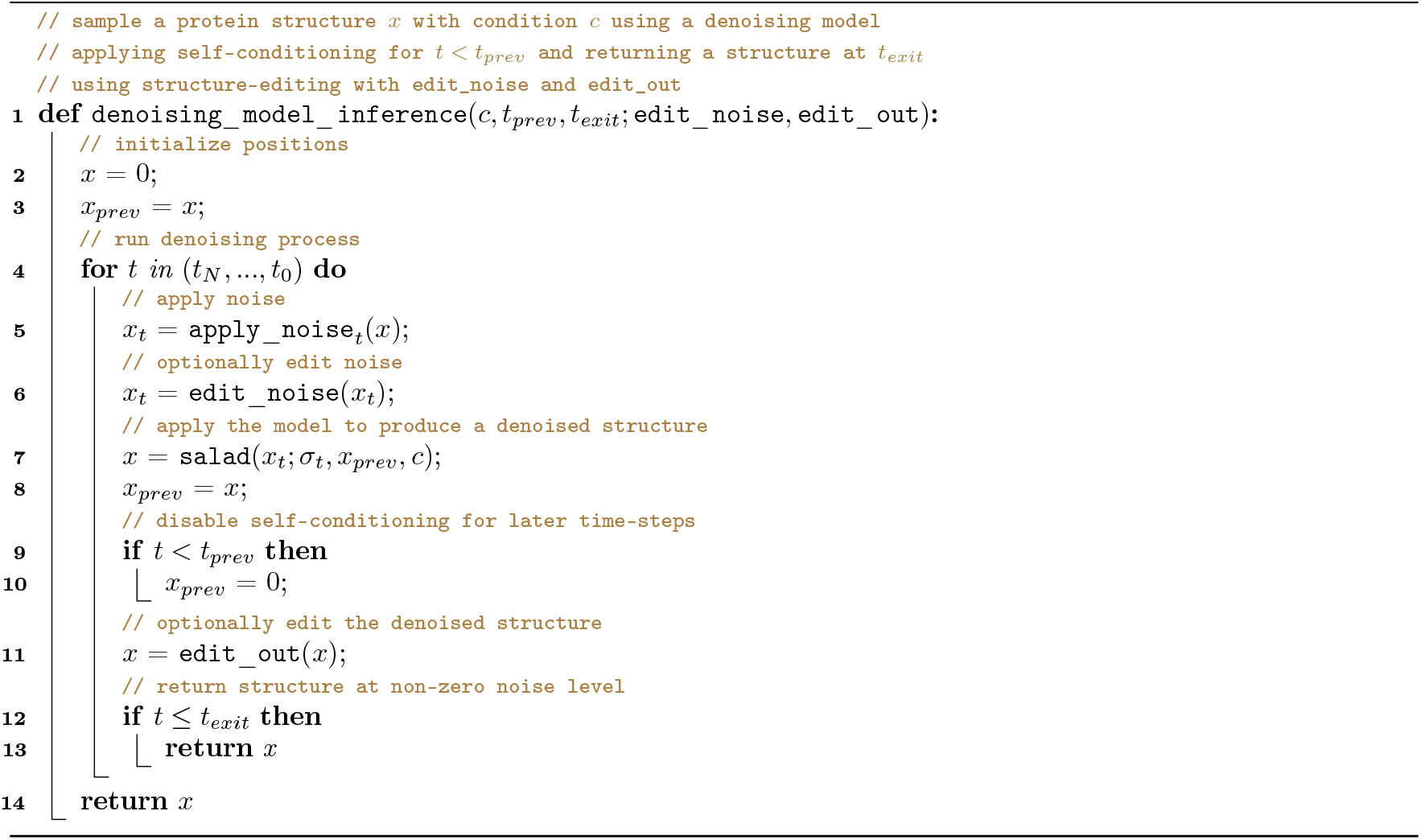

### B. Model architecture

Our sparse denoising models consist of 3 separate modules. An `Encoder` that encodes the ground-truth backbone atom positions (N, CA, C, O and idealized CB) *x*_*gt*_ and adds 15 additional pseudoatom positions for each residue to result in the denoising model input *x*, a `DenoisingModule` that receives noised positions *x*_*t*_ and is trained to reconstruct *x* and an `AADecoder`, which predicts an amino acid sequence and side chain conformations for each residue. The model is trained with self-conditioning, receiving a previously predicted structure *x*_*prev*_ and per-residue representation `local`_*prev*_ as additional inputs. All modules are based on a sparse transformer architecture [35] with pre-normalization [37].

#### Denoising module

The `DenoisingModule` consist of 6 denoising blocks based on a pre-norm transformer architecture [37]. Every block updates the per-residue representation `local`_*i*_ of size `local_size` = 256 and residue atom positions *x*. We save the trajectory of *x* values across all blocks to apply losses over the entire denoising trajectory (Sec. IV A).

Algorithm 2 shows an overview of a block in the `DenoisingModule`. We replace standard self-attention in the transformer block by a sparse version of invariant point attention (`SparseIPA`) [21]. Instead of computing the attention matrix and pair features for all amino acid pairs, we compute them for a set of precomputed neighbours. This reduces the complexity of attention from *O*(*N* ^2^) to *O*(*N* · *K*) where *K* is the number of neighbours per residue. To support conditioning on structure information, we use two sparse IPA layers. The first IPA layer operates on the current set of position features while the second operate on previous positions from self-conditioning, as well as block contact and distance conditioning information. For multi-motif models, we instead run IPA using motif information first, followed by IPA on the current position features.

Following `SparseIPA`, the per residue features `local`_*i*_ are updated using a GeLU-gated feed-forward layer with global pooling of the hidden state (`Update`, Algorithm 5). This combination of sparse attention and global mean pooling of features allows the `DenoisingModule` to learn global dependencies without having to use full *O*(*N* ^2^) attention.

##### Algorithm 2

Denoising module

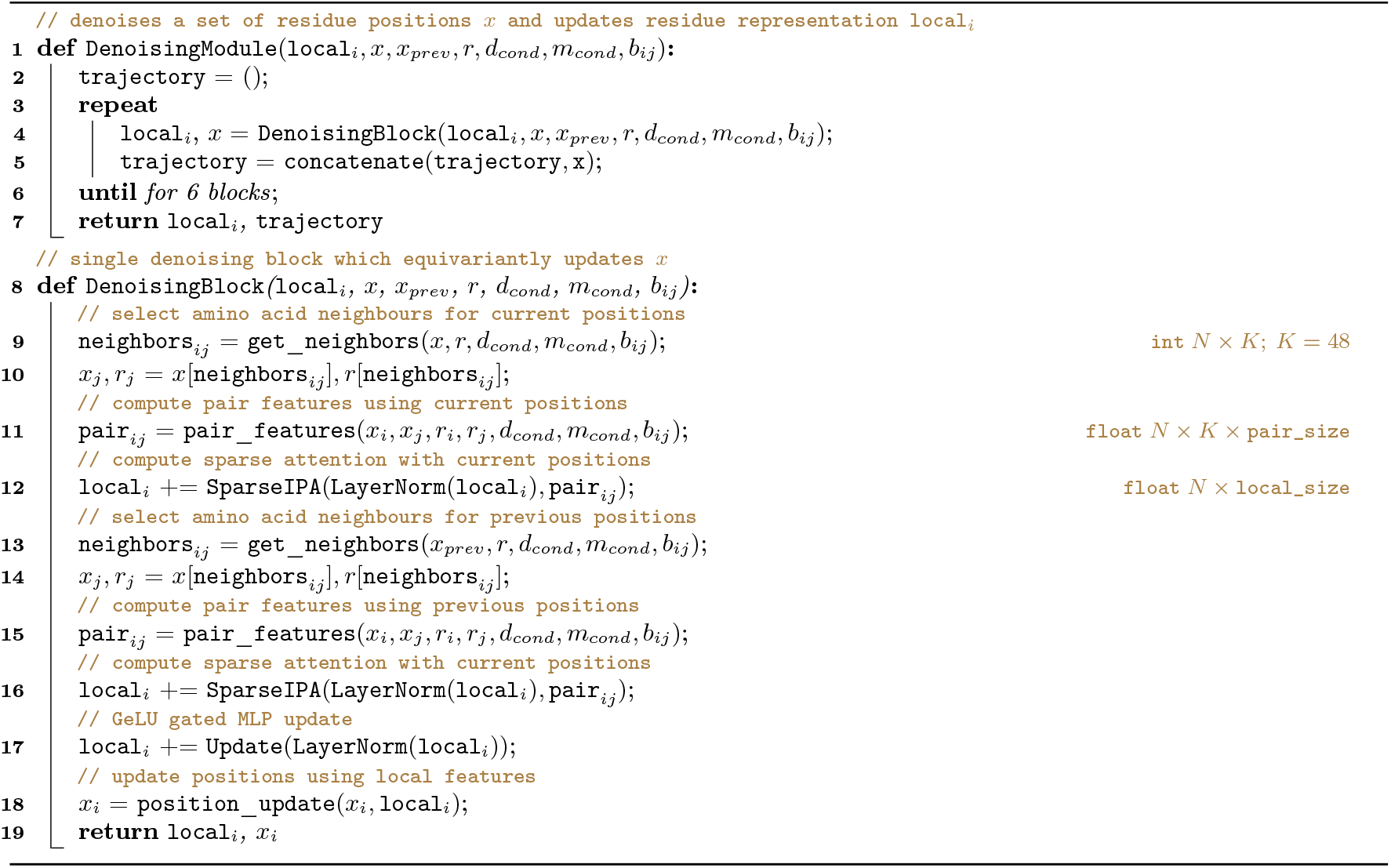

#### Neighbour selection

To compute sparse attention features, we select a set of neighbours for each residue based on their sequence and CA distances. For each residue, we choose the 16 nearest neighbours by residue index. Then we select an additional 16 neighbours by CA distance, excluding previously selected neighbours. Finally we select #`random` neighbours at random with probability 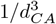 following [20] and #`cond` neighbours based on pairwise conditioning information, such as block contact conditioning [1] or pairwise distances. All default models have #`random` = 32 when computing neighbours on the current set of positions and #`random` = #`cond` = 16 when computing neighbours on self-conditioning information. This results in a total of 64 neighbours per amino acid. Multi-motif conditioned models have #`random` = 32 and do not use additional neighbours from conditioning information. It is important to note that unlike Chroma [20], a new set of neighbours is computed for each `DenoisingBlock`, as each block updates the residue positions *x*.

#### Pair features

As part of `SparseIPA` we compute amino acid pair features for each amino acid and its selected neighbours. Distances between all pairs of backbone atoms (N, CA, C, O and idealized CB) are computed for each pair and featurized using 16 Gaussian RBF [23] uniformly spaced between 0Å and 22Å. The bandwidth is set to the distance between RBF centers σ= 1.375Å. In addition to distance features, we also compute direction from the CA atom of each residue to each of its neighbour, the relative rotation between residues and the atom positions of a residue and its neighbour in local coordinates. These features are then flattened and linearly projected to `pair_size` = 64 pair features. For models with minimal pair features, we instead only compute inter-residue distances and relative rotations (Algorithm 4)

##### Algorithm 3

neighbour selection

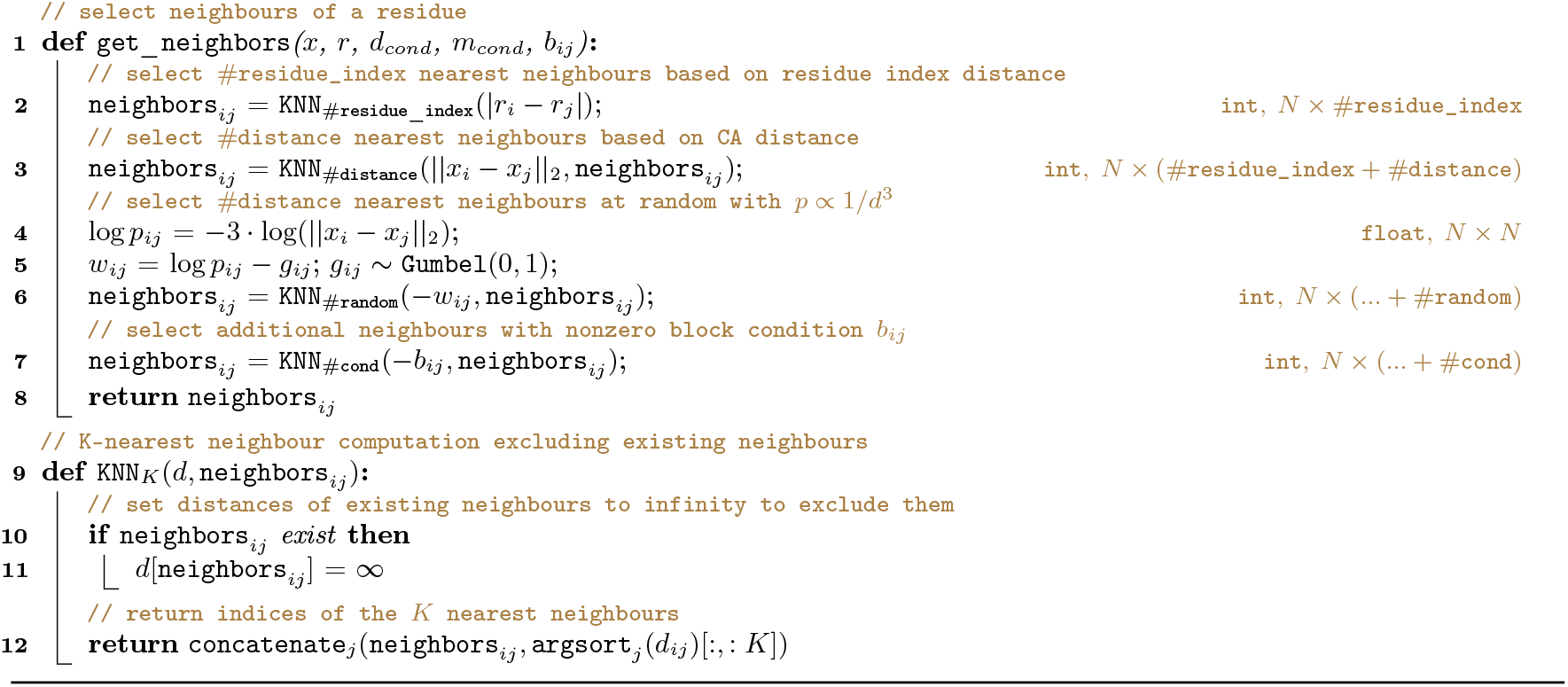

##### Algorithm 4

Residue pair features

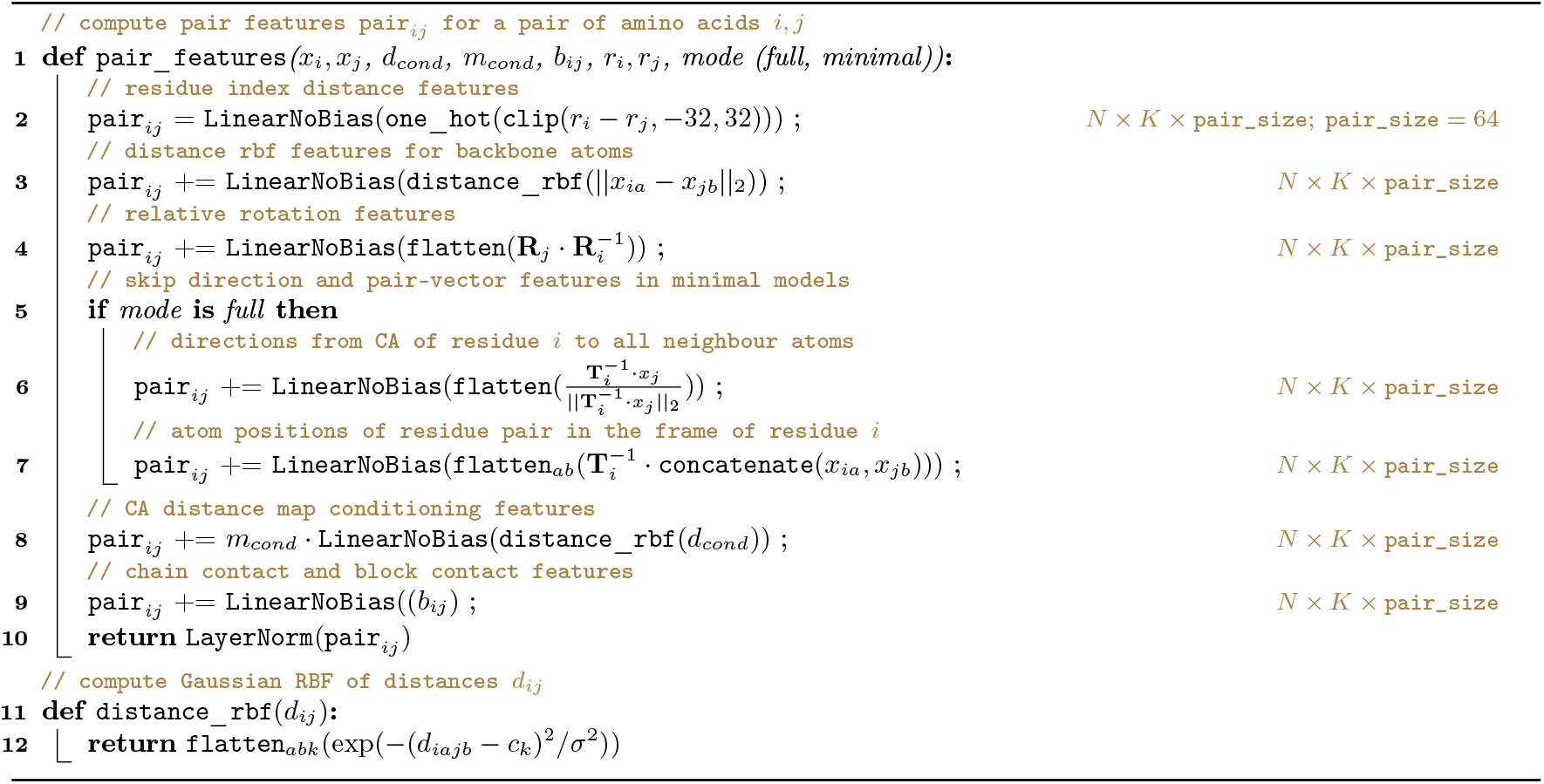

#### Update module

After applying `SparseIPA`, we update per-residue features `local`_*i*_ using a gated feed-forward layer. We first update `local`_*i*_ using the atom positions in each residue. Then, we linearly project and pool amino acid features within and across chains. Per residue, per chain and per complex features are then summed and passed through a final linear layer (Algorithm 5). Combined with `SparseIPA`, this allows the model to learn global dependencies within a protein complex without the need for full *O*(*N* ^2^) attention.

#### Equivariant position update

The final component of a `DenoisingBlock` updates the atom positions *x*_*i*_ for each residue in an equivariant manner. Per residue features `local`_*i*_ are linearly projected to a set of position updates, scaled by a unit factor of 10Å and added to the current positions *x*_*i*_ in the local frame of each residue *i*. The resulting updated positions are then transformed back into global coordinates (Algorithm 6).

##### Algorithm 5

Lightweight global update

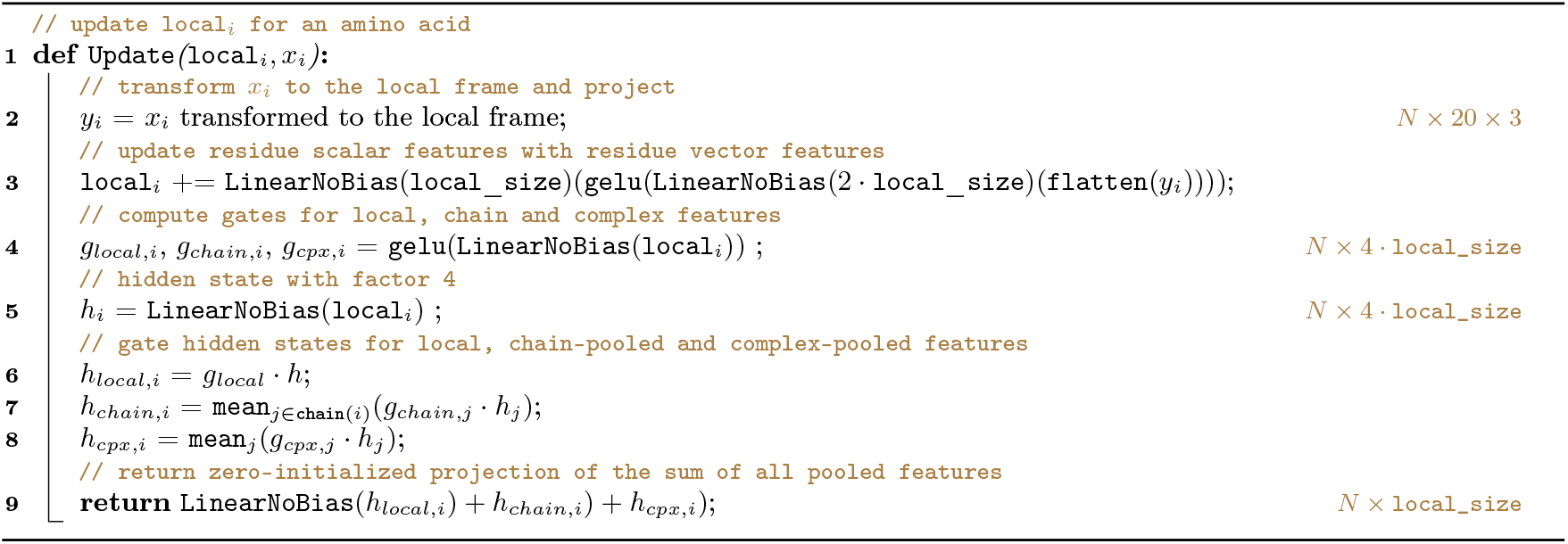

##### Algorithm 6

Position update

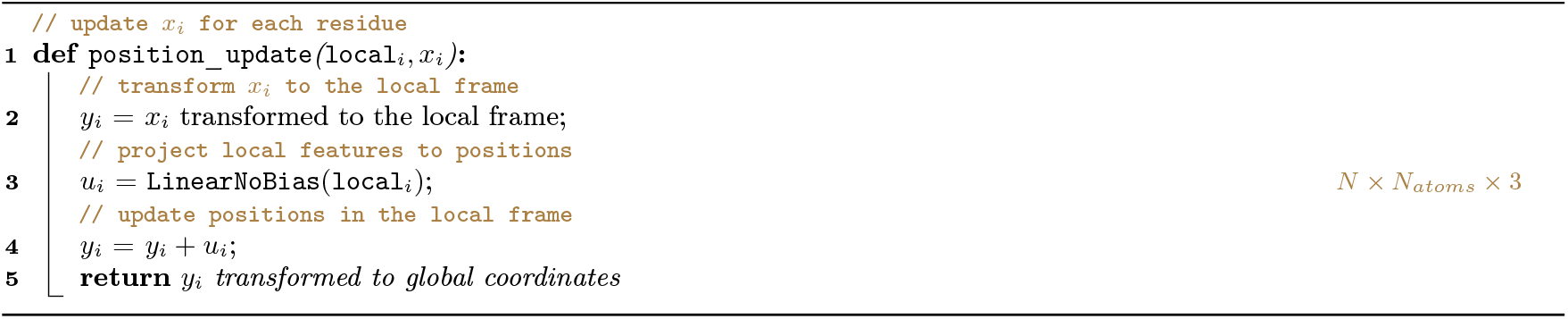

#### Structure Encoder

The `Encoder` uses a simplified version of the `DenoisingModule` and uses the same feature size as the main `DenoisingModule`. As the encoder does not change the protein backbone *x*_*gt*_, we use a precomputed set of neighbours for each residue. Each amino acid is assigned a set of 32 nearest neighbours based on CA distance. The trunk of the `Encoder` consists of two blocks of `SparseIPA` followed by a GeGLU layer [36]. After the second block of the `Encoder`, the residue representation is used to generate 15 pseudoatom positions per residue which are combined with the backbone atom positions. The resulting structure *x* is used to train the `DenoisingModule`.

#### Amino acid decoder

The `AADecoder` uses 3 blocks of the same type as the `Encoder`, together with a set of 32 nearest neighbours per amino acid computed on the denoised CA positions. In addition to denoised positions *x* and `DenoisingModule` features `local`_*i*_, the `AADecoder` also receives a partially masked amino acid sequence during training. A random fraction between 1 % and 100 % of amino acids in each training sequencs are replaced by a mask token. The `AADecoder` is then trained to predict the masked amino acids with a cross-entropy loss. This corresponds to the training objective of an autoregressive diffusion model [61].

#### Model variants

We trained denoising models for 3 different noise schedules: VP with *σ*= 10Å; VP with *σ*= *σ*(*x*_*CA*_) dependent on the standard deviation of CA atoms in the training example; and VE diffusion with *σ*∼ `LogNormal`(1.4Å, 1.6Å). For each noise schedule, we trained 3 ablated models: a model with full pair features (Sec. 12) and Fourier time embedding [29]; a model without time embedding features; a model with minimal pair features and no time embedding features.

### C. Denoising model loss

Our denoising models are trained using a combination of standard denoising and auxiliary losses. A per-block denoising loss is computed on residue (pseudo) atom positions for the output 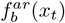 of each `DenoisingBlock`, where *r* are residues and *a* are the atoms in each residue:

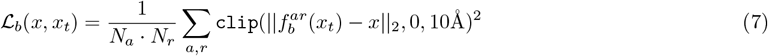

The norm ||*x*_*b*_ − *x*|| is clipped to 10 Å to stabilise training and the loss is averaged over residues *r* and (pseudo) atoms *a* in each residue. The losses for each block are then weighted together to result in a trajectory denoising loss

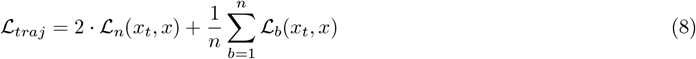

here, the final prediction is weighted by a factor of 2 to increase its importance in the final loss. This is combined with an auxiliary all-atom denoising loss using the all atom structure 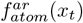 predicted by the `AADecoder`:

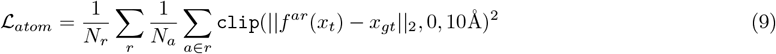

To ensure that the models learn to reproduce the relative orientations between amino acids, we also introduce a rotation denoising loss for each block following RFdiffusion [1]

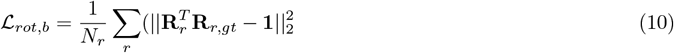

where R_*r*_, R_*r,gt*_ are the rotation matrix defined by the backbone frame of each residue in the predicted and ground-truth structures, respectively [1]. This results in a trajectory denoising loss for residue rotations

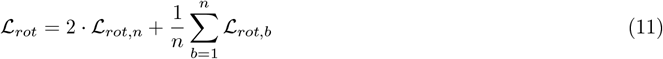

In addition to using unaligned denoising losses, we also compute a squared frame-aligned point error (FAPE) loss 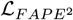 [21] over the trajectory of predictions 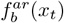 as well as a local FAPE loss on the predicted all atom structure ℒ _*local*_. Instead of computing the FAPE over all amino acid pairs, we instead compute it over the nearest 64 neighbours in the ground truth structure ℒ _*fape*_ and 16 nearest neighbours for ℒ _*local*_. As with the denoising loss, the FAPE losses are also clipped to a maximum of 10 Å. Finally, the structural losses also include AlphaFold’s structural violation loss ℒ _*viol*_ [21] to penalize clashes in denoised structures.

The models are also trained with a number of non-coordinate losses, consisting of a distogram ℒ _*dist*_ [21] and amino acid prediction ℒ _*aa*_ and secondary structure ℒ _*dssp*_ cross entropy losses:

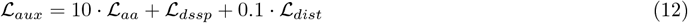

The final weighted loss of the model is then

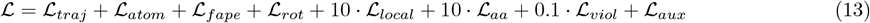

In this loss, ℒ _*viol*_ and ℒ _*local*_ are set to zero in the high-noise regime (diffusion time *t >* 0.5 for VP models; noise *σ*_*t*_ *>* 5.0Å for VE models), as the model is unlikely to learn to predict non-clashing structures at high noise levels.

### D. Structure autoencoder models

Our sparse autoencoders were implemented to have the same graph transformer architecture as the denoising models. Each autoencoder model consists of a single `Encoder` block with `SparseIPA` over 32 nearest neighbours for each residue, followed by a GeGLU layer [36]. The resulting per-residue representation `local`_*i*_ is then layer-normalized [62] and linearly projected to a latent vector *z*_*i*_ : ℝ^latent_size^ for each residue *i*. For model variants with vector quantization (VQ) [39] enabled, *z*_*i*_ is then quantized with a codebook of size 4,096.

This latent representation is decoded by a `Decoder`, which consists of 6 blocks of `SparseIPA` followed by the same `Update` and `position_update` layers used in our denoising models. The decoder is trained with 0 to 3 recycling steps for each batch, initializing the positions *x* of each recycling step with the result of the previous step *x*_*prev*_. The first recycling step starts from randomly initialized positions *x* ∼ 𝒩 (0, 1). We train models with 3 different decoder variants: an SE(3) equivariant model (EQ) using the same neighbour selection and features as used in our denoising models; an equivariant model with per-block distogram prediction (EQ+dist) and an additional `SparseIPA` layer using distogram nearest neighbours; a non-equivariant model (NEQ) directly embedding atom positions without first projecting them to residue local coordinate systems. This results in a total of 6 models trained (3 decoder variants with and without VQ).

The models are all trained with 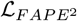 over the entire denoising trajectory and ℒ_*local*_ for the final all-atom structure prediction. Amino acid cross-entropy ℒ _*aa*_ is used as an auxiliary loss. In addition, models with per-block distogram loss are trained with distogram cross-entropy for each layer ℒ _*dist*_. This results in a combined autoencoder loss of

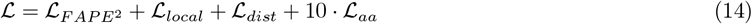

### E. Training dataset

We trained our models on a snapshot of PDB collected in October 2023 excluding any PDB entries submitted after December 31, 2020. PDB entries were then filtered for a resolution <= 4 Å. Entries containing protein chains of length < 16 were excluded from training and non-amino acid residues were removed from chains in the data set. We clustered all protein chains in the resulting data set using mmseqs2 (git commit 4f046dd) [63] with a 30% sequence identity cut-off. To train our model, we generated input micro of 1024 amino acids. Batches were constructed by repeatedly sampling structures from the data set, until the total number of amino acids reached 1024. If the total number of amino acids would exceed 1024, the batch was instead zero-padded, and the sampled structure included in the next batch. At each epoch, we sampled clusters from the data set without replacement, selecting a random chain identifier and biological assembly for each cluster. If the selected chain belonged to a complex and the entire complex fit in the current batch, we added the complex to the batch with probability 50 %. Otherwise, we added the selected chain.

### F. Model training

All denoising models were trained for 200,000 iterations on the dataset described in Section IV E with mini-batch size 1024 and 32 batches per iteration, resulting in a total of 32,768 residues per iteration. Structure autoencoder models were trained for 200,000 steps with a batch size of 16,384 residues. We used the Adam optimiser [64] with *β*_1_ = 0.9 and *β*_2_ = 0.99. The learning rate was warmed up from 0 for 1,000 steps at the start of training and then reduced to 1e-7 using cosine decay [65]. On an example machine with 8 nvidia RTX 3090 GPUs, an average training run took 3.5 days, or 672 GPU hours. Models were trained on different GPU nodes using 8 of either nvidia RTX 3090, A40 or L40s GPUs.

### G. Runtime benchmarking

We compared runtimes of salad models to RFdiffusion, Genie2, and Chroma on a single nvidia RTX 3090 GPU. We sampled 11 structures from each model using their default settings (Table I) and measured the time elapsed for each generated structure. We discarded times measured for the first generated structure to account for library initialization and model compilation and reported the average time for the remaining 10 generations.

**TABLE I.** Default model settings for runtime benchmark.

| model | # steps |
| --- | --- |
| salad | 400 |
| Genie2 | 1,000 |
| RFdiffusion | 50 |
| Chroma | 500 |

### H. Model ablation study

We selected a model architecture and sampling hyperparameters by evaluating models with and without time-embedding features as well as with full and minimal pair features (Sec. 12) on unconditional structure generation. We generate 200 backbones for proteins of size 50 to 400 amino acids for each model, using 100, 200 and 500 diffusion steps with early stopping at 80, 180 and 400 steps into the denoising process. Self-conditioning was applied until diffusion time *t*_*prev*_ = 0.8 for VP models as this was determined to yield good results in preliminary testing. For VE models, we tested self-conditioning thresholds of 0.8 and 0.99. 10 sequences were designed for each backbone using ProteinMPNN [23]. Structures were predicted using ESMfold [22]. Designability was measured as the fraction of backbones with at least one designed sequence with pLDDT > 70 and scRMSD < 2.0 Å (Extended Data Fig. 6 A, B). Models with full pair features, time-embedding and 500-step sampling were chosen for further benchmarking. *t*_*prev*_ = 0.99 was chosen for VE models.

### I. Unconditional generation benchmark

At each evaluated protein length between 50 and 1,000 amino acids, we generated 200 protein backbones using both our models and Genie2 / RFdiffusion for comparison [1, 2]. Backbones were sampled using 500 diffusion time-steps with early stopping at 400 time steps and self-conditioning turned off below the threshold diffusion time *t*_*prev*_ = 0.8 for VP models and *t*_*prev*_ = 0.99 for VE models. For models based on variance preserving (VP) noise [29] (RFdiffusion, Genie 2, our VP model), we stopped generation at proteins with length 400 amino acids, as we could already observe deteriorating designability. For each backbone, we then generated 10 amino acid sequences using ProteinMPNN with temperature 0.1 [23]. This resulted in a total of 11 sequences for our models (10 ProteinMPNN + 1 from the model itself) compared to 10 sequences for Genie2 and RFdiffusion. To fairly measure model performance and remain comparable to previous work, we restricted all computed performance measures to use the first 8 sequences generated by ProteinMPNN.

We predicted the structures of each sequence using ESMfold [22] and AlphaFold 2 [21]. For each structure prediction we measured the RMSD to the generated backbone (scRMSD) and pLDDT. Following Lin *et al*. 2024 [2], we then computed designability as the fraction of generated backbones with at least one sequence with ESMfold pLDDT > 70 and scRMSD < 2 Å. We evaluated pairwise similarities between generated backbones using TMalign (version 20220412) [66]. To compute backbone diversity for direct comparison with Genie2 and RFdiffusion [2], we randomly subsampled the set of generated structures to a size of 100 backbones. Designable structures in this subset were then clustered using single-linkage clustering on the TM-score. Backbones with TM-score > 0.6 were included in the same cluster. Diversity for all backbones (`diversity`_*all*_) was then defined as the fraction of designable clusters in all generated structures 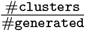 [2]. We also defined a second diversity measure as the fraction of clusters in all designable structures 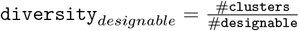 to fully separate diversity from designability. Diversity was measured on 10 samples of 100 structures each sampled from the original 200 generated structures to report median, minimum, and maximum diversity for each model.

### J. Shape initialized structure generation

We prepared letter shapes as paths in SVG format using Inkscape 1.4 (e7c3feb100, 2024-10-09) [67] then extracted the coordinates of the nodes in each path into a CSV file. To sample structures based on these shapes, we used our variance expanding (VE) model with default settings and shaped noise initialization. Instead of initialising the denoising process with noise for each residue *i*

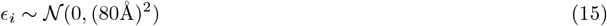

we instead centered the noise on the coordinates of nodes of the SVG path corresponding to the desired shape:

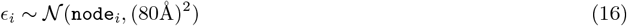

where `node`_*i*_ is the position of the node assigned to residue *i*. To generate the letter shapes described in this work, we assigned 200 consecutive residues to each node, i.e. residues 1-200 were assigned to the first node, 201-400 to the second, and so on. We then sampled 10 structures for each letter shape (S, A, L, D) starting from this noise, designed 10 sequences with ProteinMPNN at temperature 0.1 and predicted their structures with ESMfold. We then identified designable structures as in Section IV I.

### K. Domain-shaped noise

To starting noise for VE models better suited for large protein generation than normal-distributed noise, we adapted shape-initialized noise generation (Section IV J) to work with random noise centers. We sampled random starting positions for centers `center`_*i*_ according to

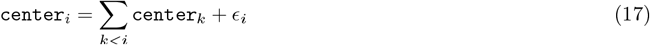

where ϵ_*i*_, ∼ 𝒩 (0, (10Å)^2^) is a normal-distributed offset. Essentially, we are sampling centers as Gaussian chains with average segment-length 10Å [20], We then enforce globularity of the chain by optimizing inter-chain distances with a harmonic restraint centered on 10Å. Optimization is done using 10 steps of gradient descent with learning rate 0.1. We then sample shape-initialized noise with 200 residues per center using these randomly generated centers. We generate fresh centers for each designed backbone.

### L. Random secondary structure conditioning

To sample a random three-state secondary structure (helix, strand, loop) of a fixed length, we first sampled a random secondary structure fraction with a maximum loop content of 50 % and arbitrary proportions of alpha helices and beta strands. We then computed the closest integer number of helix, strand and loop residues for this fraction at a fixed protein length. To arrange these residues into contiguous secondary structure elements we then heuristically determine the minimum and maximum number of helices and strands that can be generated using this number of residues (Table II). We sample a random number of helices and strands in this range and randomly assign residues to each helix and strand until we reach the previously computed number of residues for each secondary structure. These secondary structure elements are then randomly shuffled and the remaining loop residues are randomly placed in between.

**TABLE II.** Secondary structure element parameters.

| structure | min count | max count | min length |
| --- | --- | --- | --- |
| helix | $\min(1, \#\text{helix}/6)$ | $\#\text{helix}/6$ | 6 |
| strand | $\min(2, \#\text{strand}/8)$ | $\#\text{strand}/4$ | 4 |

**TABLE III.**
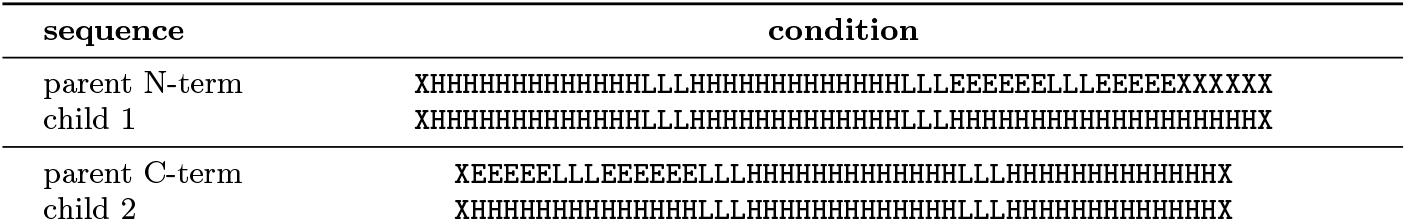
Multi-state secondary structure conditioning.

For random secondary structure sampling, we then conditioned our models on secondary structure strings generated this way, randomly replacing secondary structure elements with unknown secondary structure with a probability of 50 % per element. Additionally, the last residue in each secondary structure element was replaced with an unknown secondary structure to allow the model to decide the correct secondary structure at boundaries between secondary structure elements. Evaluation of structures generated using random secondary structures followed the procedure described in Section IV I.

### M. Synthetic dataset generation

To generate the synthetic protein dataset described in Section II D, we used our VP model with random secondary structure conditioning (Sec. IV L). We generated 50,000 backbones for random protein lengths between 50 and 256 amino acids. For each backbone, we designed 10 sequences using ProteinMPNN with temperature 0.1 [23] and predicted their structures using ESMfold [22]. We then identified successfully designed sequences with scRMSD < 2Å and pLDDT > 70. The dataset was then restricted to structures with at least one successful sequence resulting in 41,713 backbones. These backbones were then clustered using foldseek [45] with a TM-score cutoff of 0.6 and minimum coverage of query and target 0.9 using the command `foldseek easy-cluster data/pdb/ -c 0.9 –tmscore-threshold 0.6`. The coverage cut-off was chosen in this way to mostly cluster structures of similar size. This resulted in 37,661 structures which were chosen as cluster representatives and also had 1 or more successful sequence designs, corresponding to 90.3 % of designable backbones. We evaluated the percentage of novel structures relative to PDB by running foldseek [45] against PDB using TM-align and exhaustive search (`foldseek easy-search data/pdb/ fs_pdb –alignment-type 1 –format-output query,target,alntmscore,qtmscore,ttmscore,alnlen,qstart,qend,tstart,tend`, where `fs_pdb` is a precomputed copy of the PDB database downloaded using foldseek). Structures were considered novel if they had no match in the PDB with query TM-score > 0.5 (`qtmscore`).

### N. Synthetic dataset model benchmark

We trained two salad models with `default_vp` configuration on both the synthetic dataset and the PDB dataset described above (Sec. IV E) limited to sampling only single chains of length between 50 and 256 amino acids. Models were trained according to the procedure in Section IV F. We assessed the performance of both models using ESMfold structure prediction of a single sequence prediction for each generated backbone according to the procedure in Section IV I. Instead of using ProteinMPNN [23] for sequence design, we directly used the sequence defined by the argmax of the amino acid distribution predicted by each model at the final denoising step.

### O. Motif-conditional model training

To compare to Genie2 [2] we trained a separate salad model with multi-motif conditioning. The model was trained on PDB (Sec. IV E) and was given multi-motif conditioning information for each training example. Training was run for 200,000 steps according to the procedure described in Section IV F. To prepare motif conditioning information, we first partitioned each structure into contiguous segments with random lengths between 10 and 50 amino acids. Each segment was then assigned to one of 2 segment groups. Only segments within the same segment group would then be treated as a single rigid segment for the purpose of multi-motif scaffolding. Finally, segments were set as active with a probability of 50 %. Inactive segments were not used for conditioning. We then computed the CA distance map between all amino acids, together with a mask indicating amino acid pairs with active conditioning:

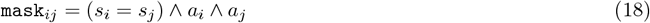

where *s*_*i*_ is the segment id of a residue and *a*_*i*_ is a boolean specifying if the segment at that residue is active.

### P. Motif conditioning using output editing

In addition to training a model for multi-motif scaffolding, we adapted the sampling process of the `default_vp` model to allow multi-motif design. At each denoising step, we align the motifs to its corresponding residues in the denoised structure. We then replace the coordinates of those residues with the coordinates of the motif (Algorithm 7). Sampling structures in this way guarantees that the motif will be incorporated into the resulting backbone.

#### Algorithm 7

Motif editing denoising process

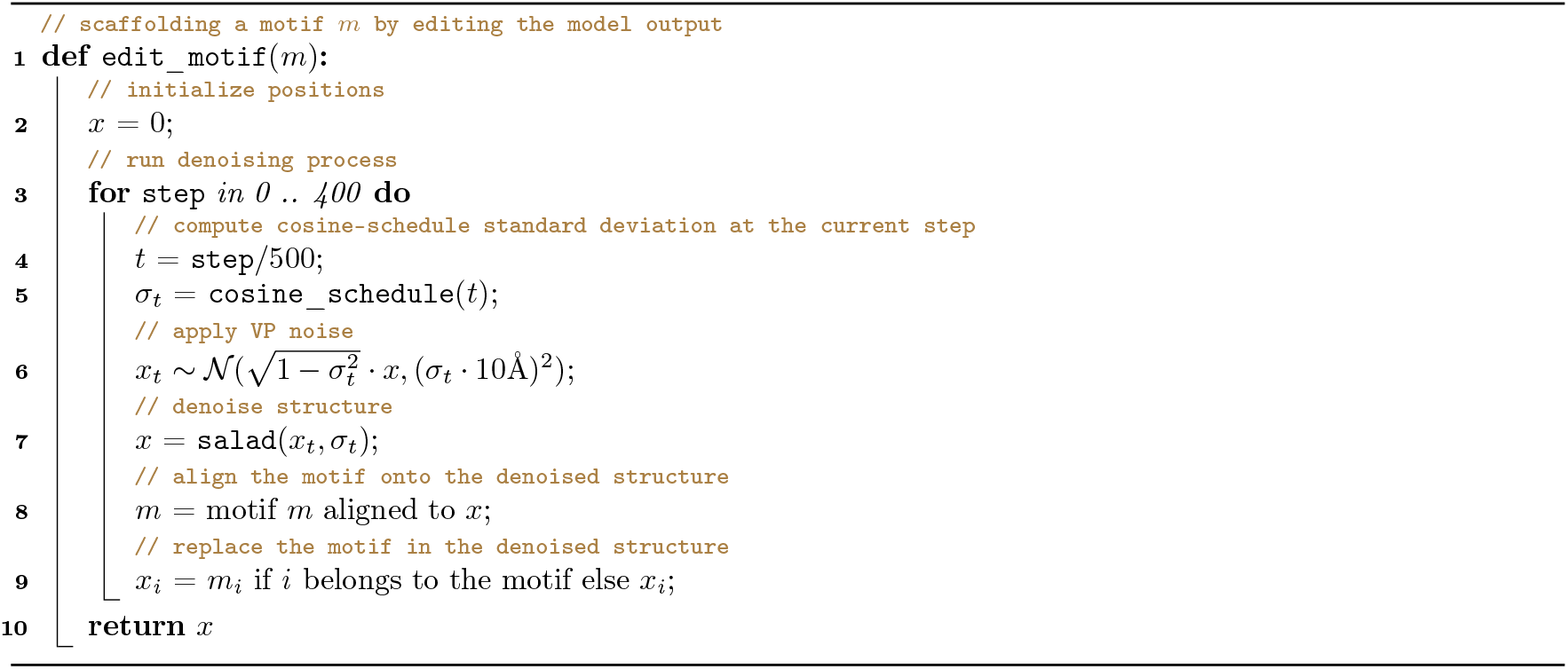

### Q. Motif benchmark

Following Lin *et al*. we generated 1,000 structures using motif-conditioning and motif-editing for each single-motif scaffolding task defined in [1] and additional multi-motif scaffolding task specified in [2]. We then designed sequences for each backbone using ProteinMPNN and evaluated designability using ESMfold (see Sec. IV I). In addition, we computed the CA and full-backbone (N, CA, C, O) RMSD between the predicted structures and the input motif. Successful designs were selected using a backbone RMSD cutoff of 1 Å and clustered using single-linkage clustering at a TM-score threshold of 0.6 to identify the number of unique successes. For comparison, we also computed the number of unique successes based on RMSD-CA. We compared these results to results published for Genie2 and RFdiffusion by Lin *et al*. using the same evaluation strategy [2].

### R. Symmetry editing

To generate symmetric repeat proteins according to a given symmetry group, a representative subunit structure was generated by aligning all subunits of a repeat protein and averaging their positions. Subunits were aligned using the action of the symmetry group. For a cyclic groups, consecutive subunits were all rotated around the symmetry axis onto a single subunit. For a screw (helical) symmetry group, consecutive subunits are first centered along the screw axis then rotated onto a single subunit around the axis. We can then position the center of mass of the subunit at a specified radius *R* from the symmetry axis to generate structures with a specified radius. The resulting representative structure is then replicated using the group action. This process is described in detail in Algorithm 8 for a group *G* with a single generator *g*.

#### Algorithm 8

Symmetry-group editing

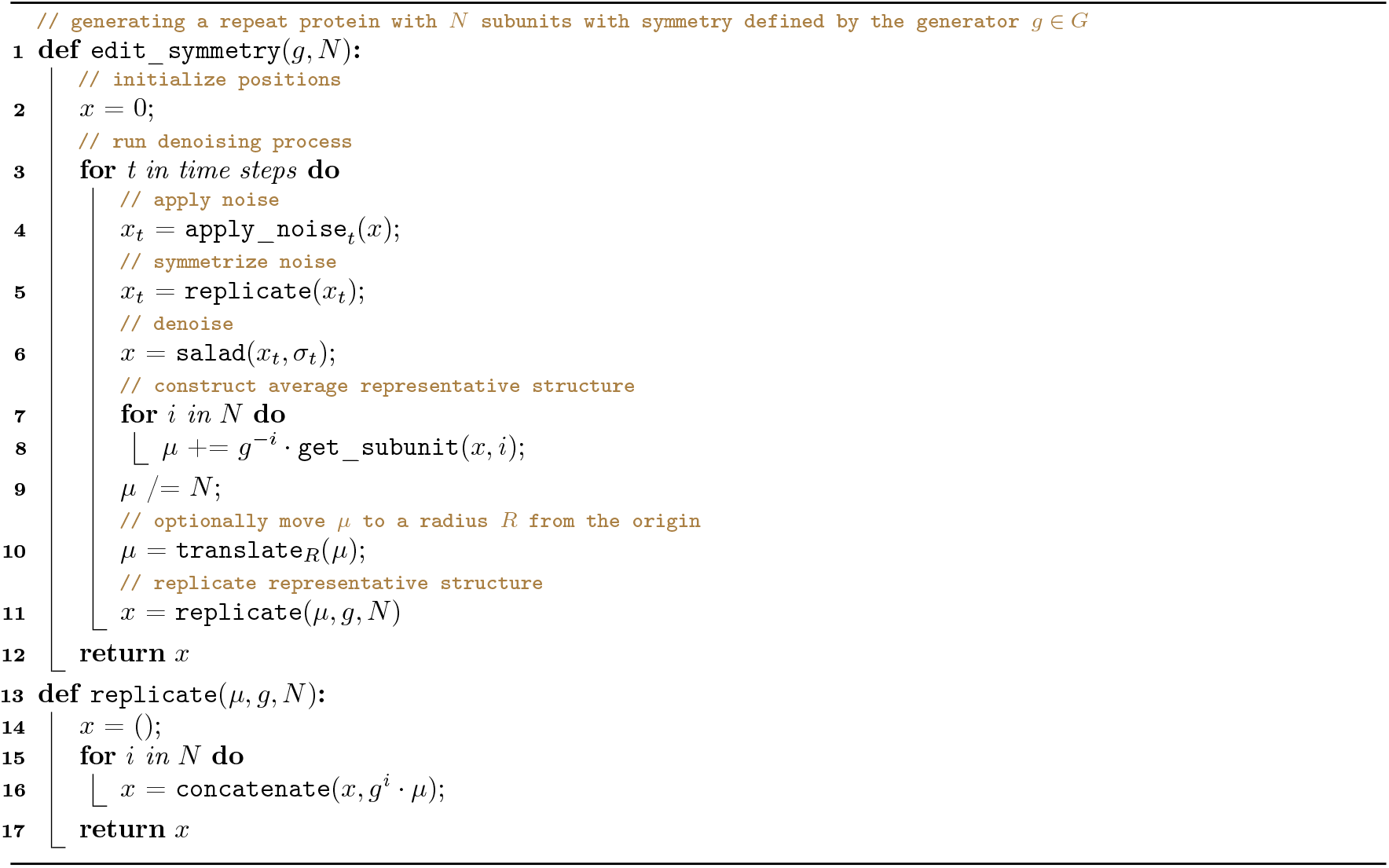

### S. Symmetry benchmark

We generated symmetric repeat proteins with subunits of length 50 and 100 for cyclic symmetry groups C3, C4 and C5 with variable radius using VP diffusion as well as C3 to C7 with radii from 10 to 14 Å using VE diffusion. In addition, we generated screw-symmetric designs with 2 to 3 repeat subunits for various angles and radii. For each design class, we generated 20 symmetrized backbones and designed 10 sequences using ProteinMPNN with temperature 0.1 [23]. We evaluated designability using ESMfold for all designs. To verify that the designed screw-symmetric proteins would be predicted to fold with more repeats added, we used [48] to verify the structure of 9-subunit repeats for a subset of designs.

### T. Multi-state output editing

Multi-state outputs were generated by running one independent diffusion process per state and editing the denoised output structures to fix shared substructures across states. To fix a set of residues across states, we aligned the fixed residue positions, optionally averaged them and copied the result back to each state. Repeating this procedure for each denoising step ensures that the fixed residues will have highly similar positions in the final generated structures. Algorithm 9 describes the editing process for a 2-state design process with a set of fixed amino acids {*m*}.

#### Algorithm 9

Multi-state editing

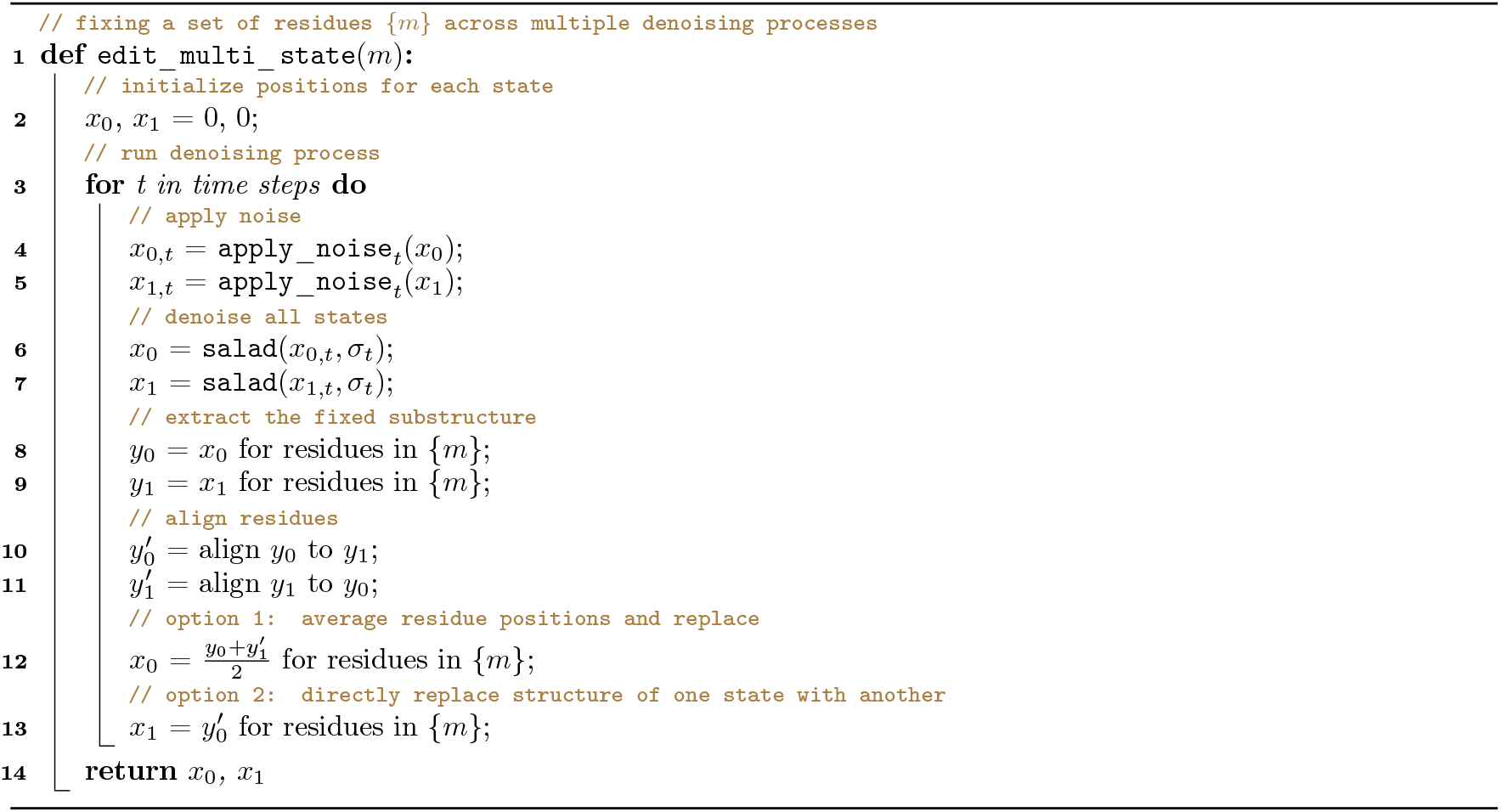

### U. Multi-state design benchmark

We generated designs for the multi-state design problem described by Lisanza *et al*. 2024 [13] (Table. 3). For each design, 3 backbones (parent, child 1, child 2) were generated with secondary structure conditioning according to using the editing strategy from Section IV T. Editing was performed with two different conditions: either the structure of the terminal helices (unconstrained), or the structure of all helices (constrained) shared between parent and children was fixed using output editing. 1,000 designs were generated per condition. We used ProteinMPNN [23] with temperature 0.1 to generate 10 sequences for each set of backbones, fixing amino acid identities across the parent and child sequences. To allow for comparison to the results presented in [13], AlphaFold2 [21] was used to determine designability. We used the cutoffs for success (scRMSD < 3 Å and pLDDT > 75) reported in [13].

## V. DATA AVAILABILITY

Generated protein structures and ESMfold-based scores and structure predictions from this work have been deposited on Zenodo. The corresponding Zenodo repository also contains the parameters of models used in this study and a snapshot of the source code that was used to generate them. The source code on [Zenodo](https://zenodo.org/records/14711580) and on GitHub contains instructions and scripts to reconstruct the datasets used for training in this study, as well as the training scripts used to produce the model parameters.

## VI. CODE AVAILABILITY

The code for all models described in this work is available under an Apache 2.0 license at github.com/mjendrusch/salad. Parameters for those models are available under a CC BY 4.0 license at github.com/mjendrusch/salad and Zenodo. The code for the AF2 and ESMfold-based benchmarking is available under an Apache 2.0 license at github.com/mjendrusch/novobench.

## VII. ACKNOWLEDGEMENTS

We thank Dr. Kashif Sadiq and the Korbel group for helpful discussions. We thank EMBL Heidelberg for generously providing access to their HPC infrastructure [68]. This work was funded by core funding from the European Molecular Biology Laboratory to the Korbel group.

## VIII. CONTRIBUTIONS

M.J. and J.O.K. conceptualized the study. M.J. conceptualized and implemented the software. M.J. designed, implemented and conducted the experiments. M.J. and J.O.K. analyzed the data. M.J. and J.O.K. wrote the initial draft of the manuscript.

## Appendix A

## Extended Data

**EXTENDED DATA FIG. 1.**
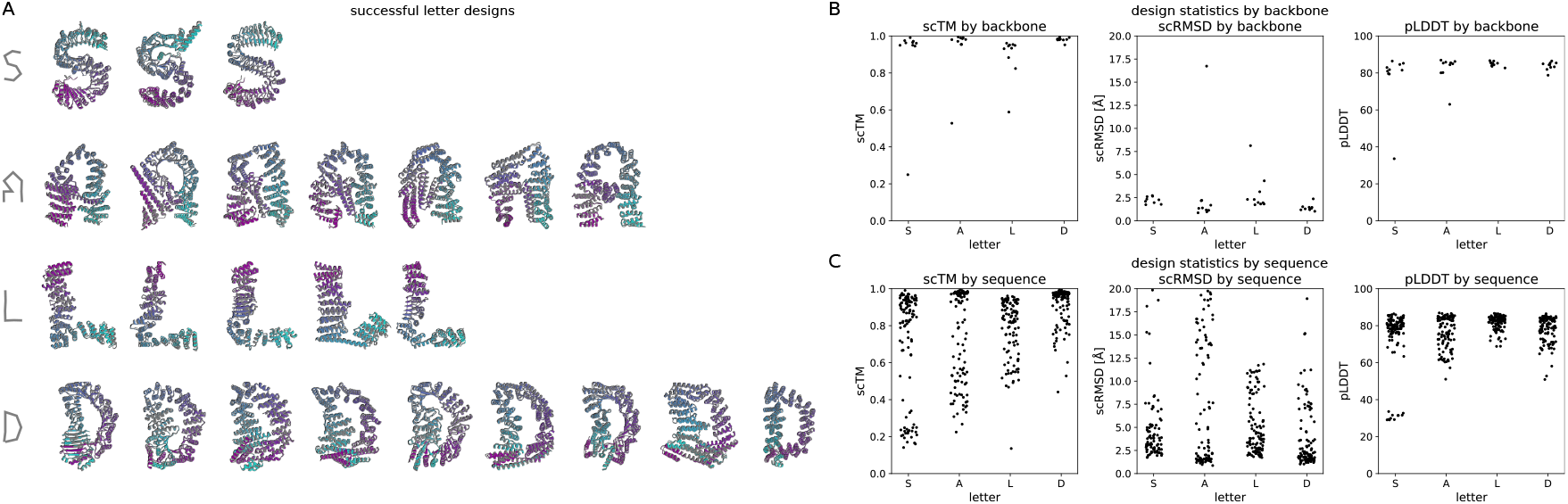
Letter shape generation. (A) Successfully designed letter shapes for letters (S, A, L, D). The designed structure (grey) is overlaid with the best ESMfold prediction (coloured by residue index) out of 10 ProteinMPNN sequences [23]. (B, C) scRMSD, scTM and pLDDT for each backbone (B) or ProteinMPNN sequence (C) across all letter designs.

**EXTENDED DATA FIG. 2.**
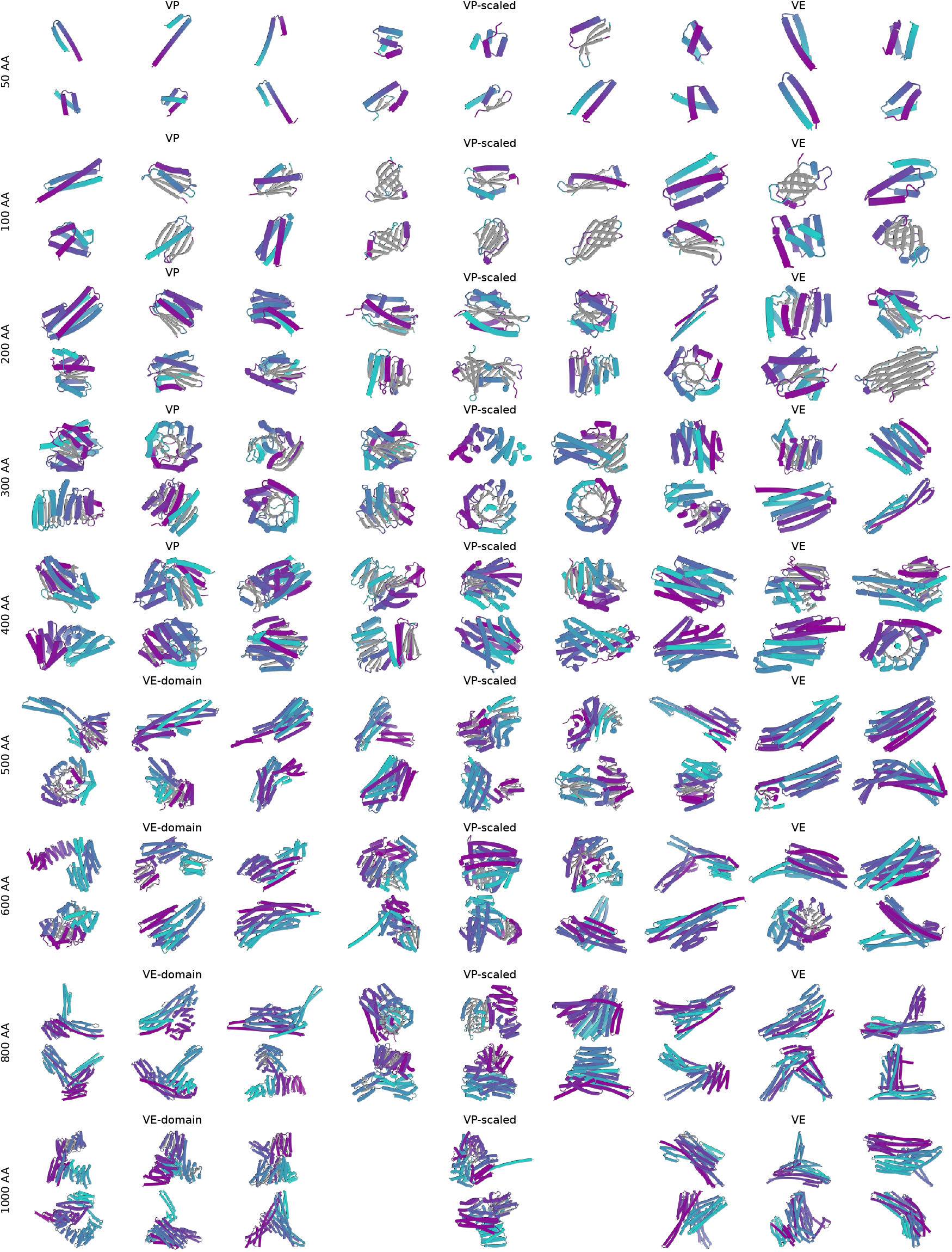
Example unconditional monomers. Monomers of length 50, 100, 200, 300, 400, 500, 600, 800 and 1,000 amino acids, using VP, VP-scaled and VE models with and without domain-like noise. Designs shown are the first up to 6 designs with scRMSD < 2.0 Å and pLDDT > 70 for each length and model type. Structures are coloured by residue index (N-terminus: purple, C-terminus: turquoise) and beta sheets are coloured in grey.

**EXTENDED DATA FIG. 3.**
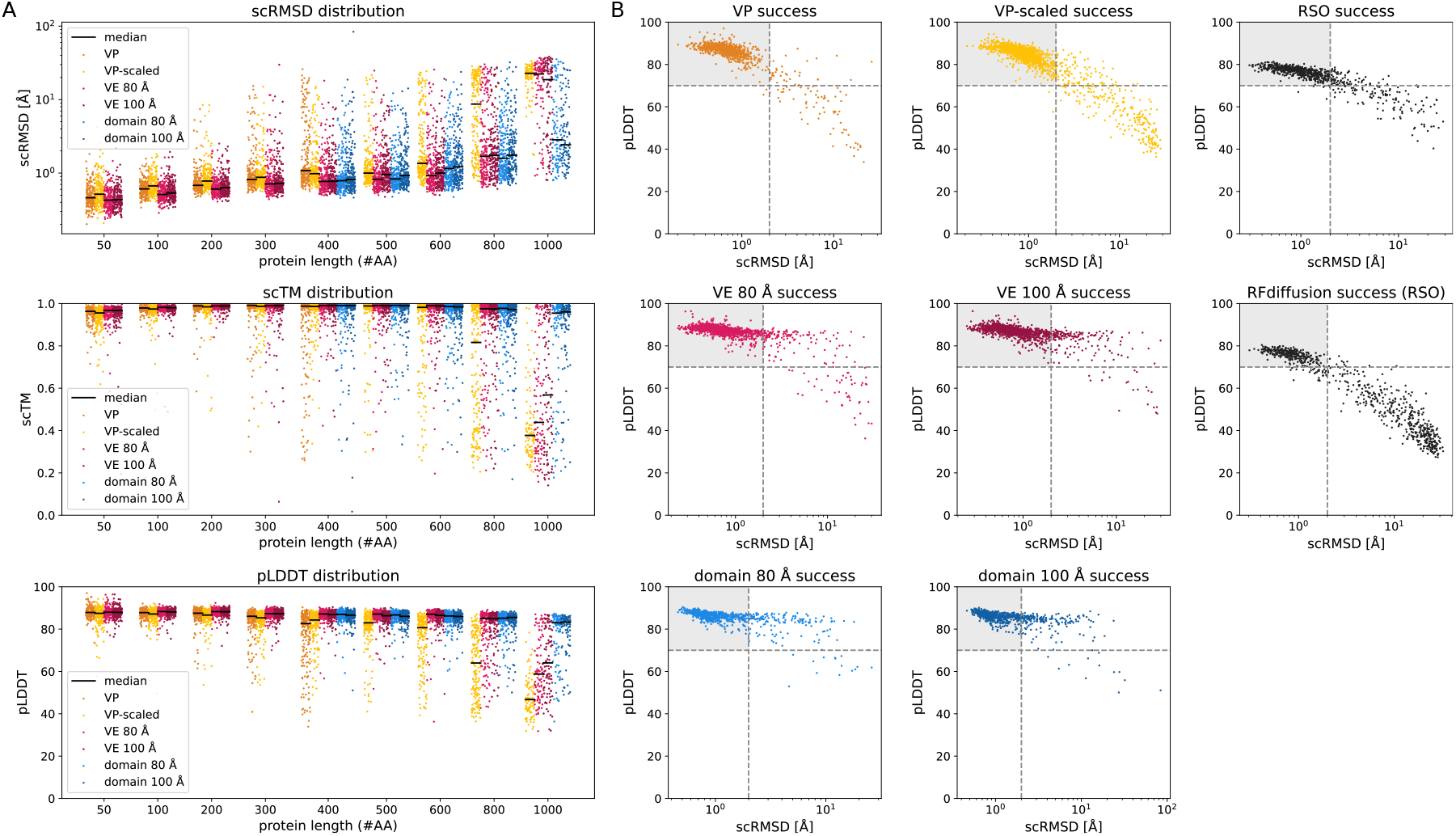
Unconditional monomer designability statistics. (A) Distributions of scRMSD, scTM between generated and ESMfold-predicted structures as well as ESMfold pLDDT for each backbone, taking the best value over 8 ProteinMPNN sequences for each backbone. Each point corresponds to a backbone. Points are coloured and grouped by model type and noise schedule. The solid black lines indicate the median value for each model. (B) Joint distributions of ESMfold scRMSD and pLDDT over all structures generated with `salad` models with results for RSO and RFdiffusion shown from comparison (results were obtained from [19]). Scatterplots are coloured as in (A). The dashed lines indicate the success cutoff values of scRMSD < 2.0 Å and pLDDT > 70 and the shaded area corresponds to the area of successful designs.

**EXTENDED DATA FIG. 4.**
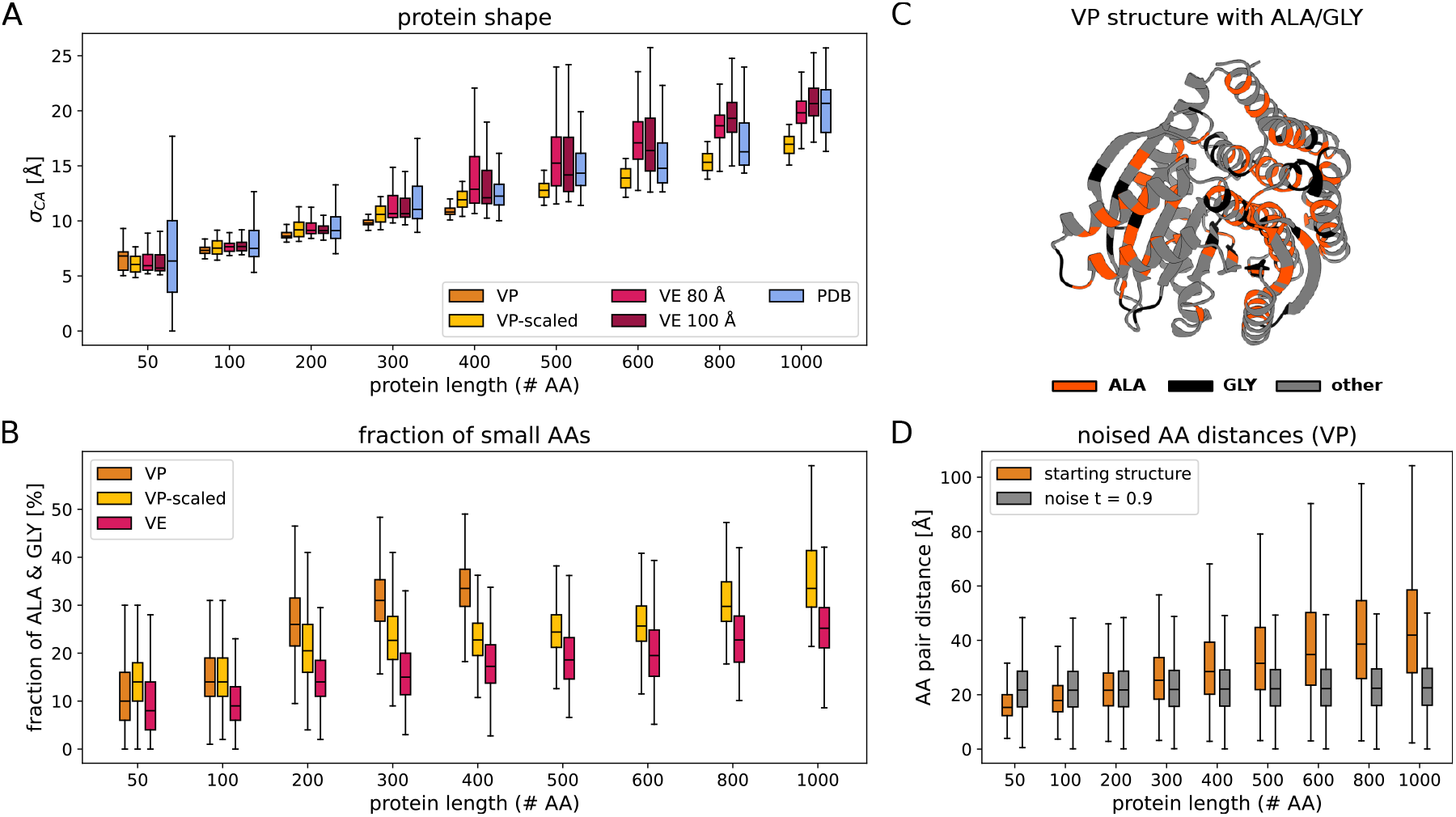
Impacts of VP noise on protein shape and sequence composition. (A) Boxplot of CA atom position standard deviations for protein structures in the PDB and backbones generated using different noise schedules. *σ*_*CA*_ distributions are shown for protein lengths between 50 and 1,000 amino acids. VP noise consistently produces highly-compact backbones with low *σ*_*CA*_, while VP-scaled and VE noise result in higher *σ*_*CA*_ structures more closely matching the *σ*_*CA*_ of proteins in the PDB. (B) Boxplots of the fraction of alanine (ALA) and glycine (GLY) residues in sequence designs for backbones generated using VP, VP-scaled and VE diffusion. A high proportion of ALA and GLY residues in a designed sequence coinsides with the presence of tightly-packed secondary structure elements with no space for more bulky amino acids. The fraction of small amino acids increases with protein length for all models but shows a particularly pronounced increase for structures generated using VP noise. In comparison, VP-scaled and VE models show a slower increase and an overall lower fraction of ALA and GLY residues. (C) Example structure of a 400 amino acids backbone generated using VP noise. ALA (GLY) residues are marked in red (black). (D) Boxplot of amino acid pair CA distances for proteins of size 50 to 400. Distributions of distances are shown for noise-free structures (orange) and noised structures at diffusion time *t* = 0.9. At low protein lengths (50 - 200), a VP model needs to decrease CA distances to denoise the structure, whereas at high protein lengths (>= 300 amino acids) it needs to increase CA distances to arrive at the denoised structure. A VP model mostly trained on smaller proteins will therefore likely develop a bias for lower CA distances resulting in overly compact, undesignable structures. (A, B, D) For all box plots, the center line indicates the median, box boundaries the 1st and 3rd quartiles and whiskers 1.5× the inter-quartile range from the box.

**EXTENDED DATA FIG. 5.**
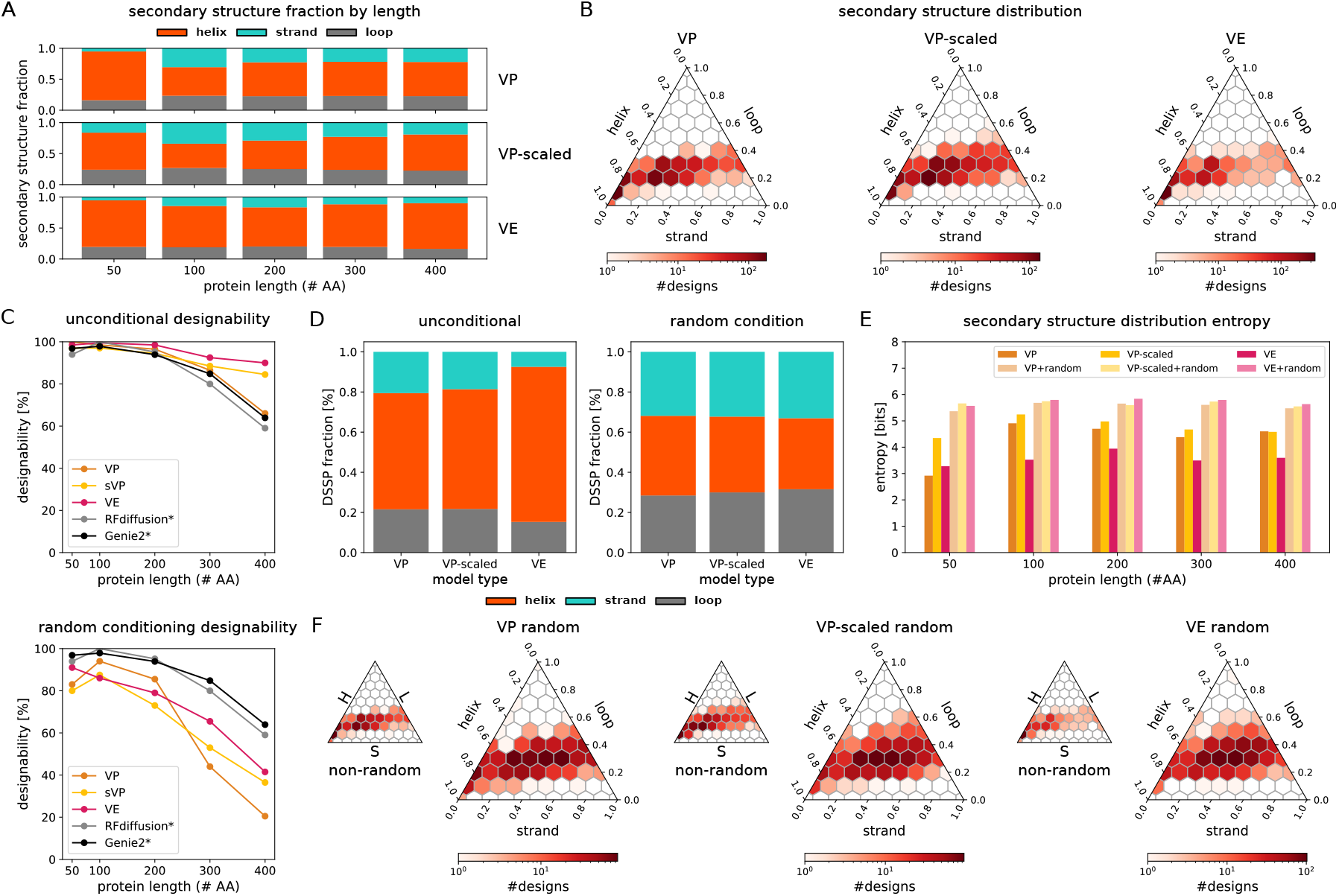
Effects of random secondary structure conditioning on diversity and designability. (A) Secondary structure fraction for protein structures between 50 and 400 amino acids generated using unconditional sampling from `salad` VP, VP-scaled and VE models. (B) Ternary plots of the distribution of secondary structure contents in generated backbones of length 50 to 400 amino acids using `salad` VP, VP-scaled and VE. (C) Designability of `salad` VP, VP-scaled (sVP) and VE generated structures for proteins of length 50 to 400 amino acids compared to results for Genie2 and RFdiffusion obtained from [2]. (top) designability for unconditional generation; (bottom) designability for random conditioning for `salad` models compared to unconditional generation for RFdiffusion / Genie2. (D) Overall secondary structure distribution for unconditional (left) and randomly conditioned (right) generations for proteins of length 50 to 400 amino acids using `salad` VP, VP-scaled and VE models. (E) Bar graph of binned secondary structure distribution entropy for unconditional and randomly conditioned generations of length 50 to 400 amino acids. The binned secondary structure distributions were constructed by subdividing the range of helix and strand percentages into 20×20 bins of equal size. (F) Ternary plots of the distribution of secondary structure content for all generated structures between 50 and 400 amino acids using `salad` VP, VP-scaled and VE models with unconditional sampling (inset) and random conditioning.

**EXTENDED DATA TABLE I.** Diffusion model hyperparameters and parameter counts.

| model | sparse | # layers | # AA features | # pair features | # parameters |
| --- | --- | --- | --- | --- | --- |
| **salad** (Encoder + Diffusion + AADecoder) | yes | 2 + 6 + 3 | 128 | 64 | **11.9M** |
| **salad** (Diffusion) | yes | 6 | 128 | 64 | **8.4M** |
| Chroma | yes | 12 | 512 | 256 | 18.5M |
| Genie 2 | no | 8 | 384 | 128 | <u>15.7M</u> |
| RFdiffusion | no | 36 | 256 | 128 | 59.8M |

**EXTENDED DATA FIG. 6.**
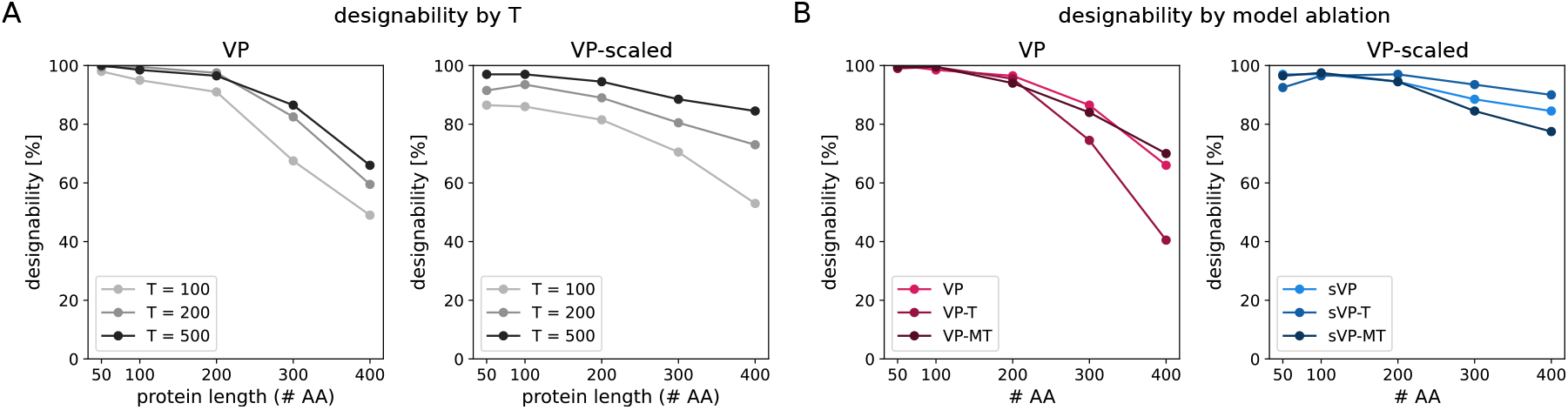
Impact of sampling steps and model architecture on designability. (A) ESMfold designability for 8 ProteinMPNN sequences per backbone of length 50 to 400 amino acids for `salad` VP and VP-scaled models at 100, 200 or 500 denoising steps. (B) ESMfold designability as in (A) for `salad` VP and VP-scaled (sVP) models at 500 denoising steps, as well as variants without diffusion time embedding (-T) and additionally using only minimal distance and orientation pair features (-MT).

**EXTENDED DATA FIG. 7.**
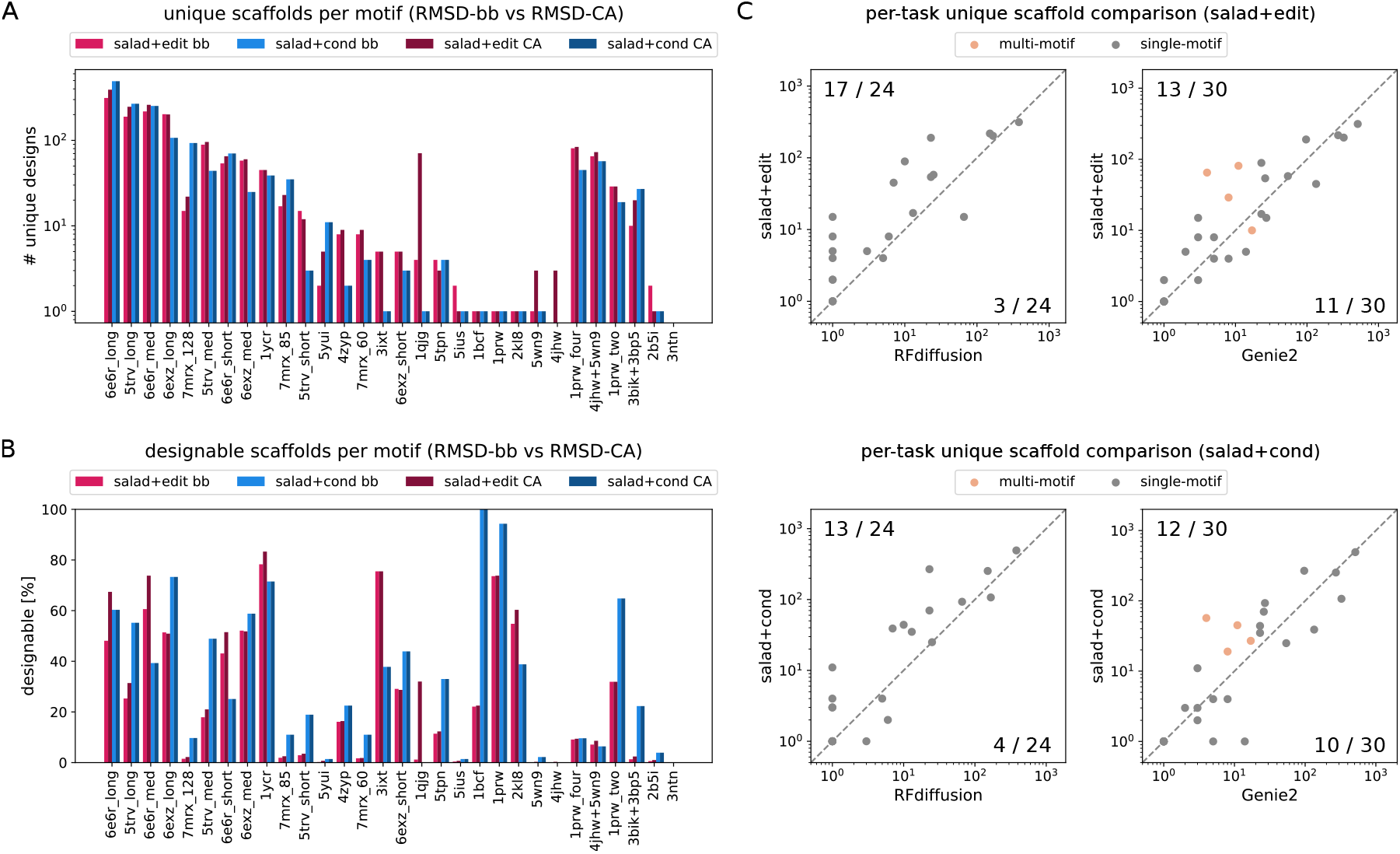
Motif scaffolding performance. (A) Number of unique scaffolds per motif-scaffolding problem using backbone atom (N, CA, C, O) RMSD (bb) and CA RMSD (CA) as a threshold for success. Results are shown for both structure-editing (`salad+edit`) and motif conditioned models (`salad+cond`). (B) Percentage of successful designs for each motif-scaffolding problem. (C) Scatter plots comparing the number of unique successful scaffolds for all single and multi-motif scaffolding tasks between `salad` models and state-of-the-art diffusion models (Genie2, RFdiffusion). The x and y axes show the number of unique scaffolds for each model. Points on the dashed line correspond to motifs with an equal number of designs for both methods. For points above the line, `salad` is better; for points below the line RFdiffusion/Genie2 is better. The number of points above and below the line is listed in the upper left and lower right corners.

**EXTENDED DATA FIG. 8.**
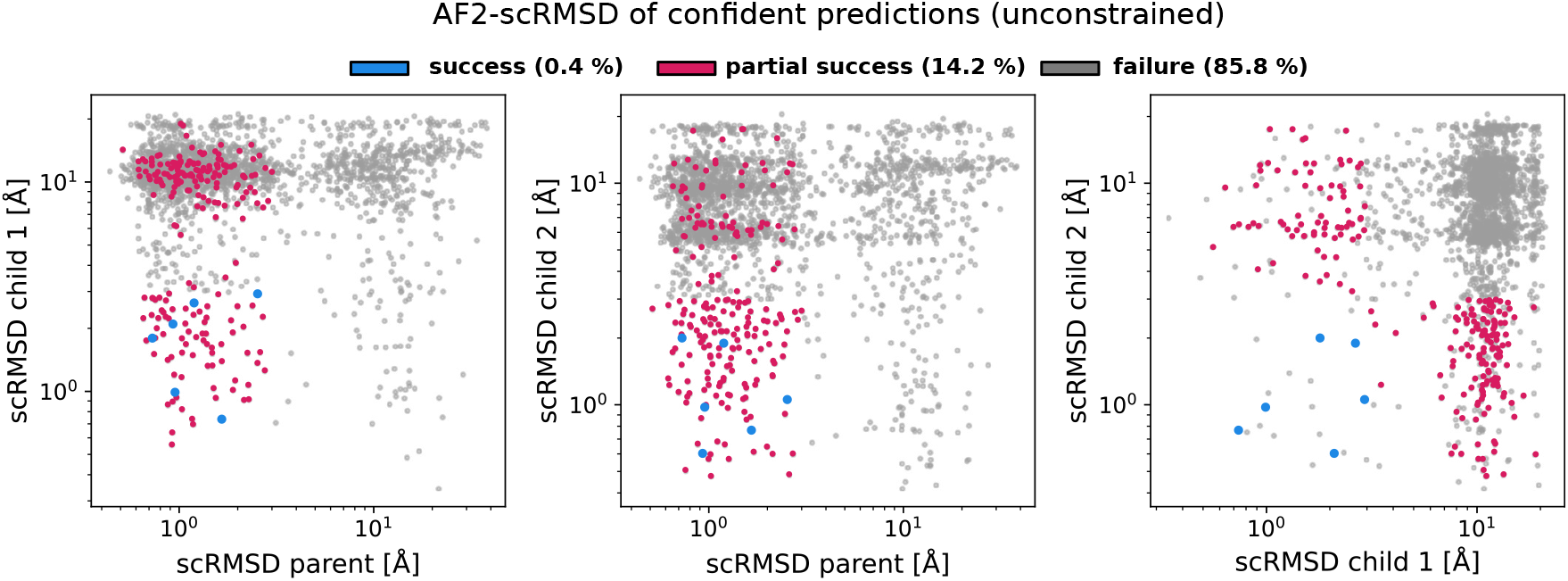
Design success for unconstrained multi-state design. Scatter plots of scRMSD for parent and child designs generated only with secondary structure conditioning, without fixing parts of the structure across denoising processes. Only designed sequences with AF2-pLDDT > 75 are shown. Successful designs (0.4 % of backbones) are shown in blue, partially successful designs (14.2 %) in red and failed designs (85.8 %) in grey.

